# Herbicolin A, an antifungal lipopeptide produced by *Pantoea agglomerans* APC 4211 is a promising biocontrol agent against food spoilage fungi

**DOI:** 10.64898/2026.05.21.726617

**Authors:** Eleni Kamilari, Paula M. O’Connor, Jerry Reen, Promi Das, Aiswariya Deliephan, Daragh Hill, Oxana Fursenko, Jonathan Weise, Ay Sha Naomi Moore, Colin Hill, Catherine Stanton, Paul R. Ross

**Author notes:** Corresponding author and email address Ross R. P.

## Abstract

Fungal contamination of food with yeast and moulds is associated with major economic losses due to spoilage and also poses health risks in the form of mycotoxin production. The strain *Pantoea agglomerans* APC 4211 isolated from leaves of *Ilex aquifolium* (holly tree) has broad spectrum antifungal activity against a variety of food spoilage fungi. Genomic analysis of the strain confirmed the presence of biosynthetic gene clusters potentially encoding for the enzymatic machinery required for the production of the antifungal lipopeptide herbicolin A. Matrix-assisted laser desorption ionization-time of flight mass spectrometry (MALDI-TOF MS) analysis of the cell-free supernatant (CFS) confirmed the presence of molecular masses corresponding to herbicolin A (1300.8 Da), and herbicolin B (1138 Da). Purified herbicolin A has desirable properties for biotechnological applications, including potent antifungal activity against a range of spoilage fungi, thermal stability and resistance to proteases. Herbicolin A has low cytotoxicity against epithelial cell lines and has minimum inhibitory concentrations (MICs) lower than those of some commercial antifungal drugs (0.2 – 2.5 µg/ml). In a model dairy system (10% skim milk), herbicolin A demonstrated excellent solubility and stability, effectively eliminating *Aspergillus niger* and *Penicillium notatum* at a concentration of 5 µg/mL. In conclusion, herbicolin A is a potent, naturally occurring antifungal agent with the potential to be applied as a biopreservative in food systems, providing a safe, clean-label, and efficient compound for synthetic preservatives replacement.

**Highlights:**

- Herbicolin A has a strong potential as a natural preservative for food protection
- Herbicolin A shows lower MICs than several antifungal agents
- Herbicolin A is stable under heat and resistant to proteolytic degradation
- Herbicolin A has strong solubility and stability in a model dairy system
- Herbicolin A indicates low cytotoxicity against epithelial cell lines

**Data summary:** The authors confirm all supporting data, code and protocols have been provided within the article or through supplementary data files.

## 1. Introduction

Microbial contamination is responsible for 1.3 billion tons of consumable food being wasted annually (Gustavsson et al., 2011; Karanth et al., 2023). Plant-associated pathogenic fungi are responsible for crop losses of up to 30%, an amount that would be sufficient to provide food for 600 million people (Avery et al., 2019). Dairy products are among the highest affected categories as a quarter are lost or wasted annually (Martin et al., 2021). Furthermore, fungal infections are a causative factor of about 11.5 million severe and 1.5 million deadly infections annually (Zhan and Liu, 2017). These numbers are estimated to increase significantly in the years ahead, owing to the rise of antibiotic resistance in pathogenic fungi. Mycotoxins, toxic metabolites produced by fungi, contribute to significant economic losses globally (around $932 million in the US) (Whittaker et al., 2020). In addition, they can lead to severe health issues, such as immune toxicity, nephrotoxicity, hepatotoxicity, growth interruptions, and carcinogenicity even when only consumed in low doses (Liu et al., 2020). Specifically, aflatoxins, alterotoxins, ochratoxins, fumonisins, trichothecene mycotoxins, and deoxynivalenol, can contribute to acute or chronic toxicity. Most mycotoxin-producing species include *Aspergillus* sp., *Penicillium* sp., *Fusarium* sp., *Candida sp*., and *Claviceps* sp. (Hathout and Aly, 2014; Schaarschmidt and Fauhl-Hassek, 2018). Furthermore, 4% of the world population exhibits allergic reactions to fungi (Denning et al., 2014). Taken together, infection and spoilage by fungi represents a major challenge to the safety and sustainability of the human and animal food supply.

To prevent microbial growth and subsequent food deterioration, the food industry uses specific chemicals, including sorbic acid, potassium sorbate, sodium diacetate, acetic acid, sodium benzoate, and calcium and sodium propionate for the purposes of food protection and safety (Liu et al., 2020). However, the detrimental effects of excessive usage of these chemical preservatives on human health and the sensorial characteristics of certain foods point to a need for the discovery and development of safe, effective solutions for food biopreservation (Mavani et al., 2024). The rising consumer demand for clean-label alternatives has prompted the necessity to consider natural replacements. Antimicrobial peptides (AMPs) represent strong candidates for shelf-life extension and preventing spoilage due to fungal contamination of food products (de la Lastra et al., 2025). Herbicolins A and B are non-ribosomally synthesized peptides (NRPs), isolated from some bacterial strains of *Pantoea agglomerans* (formerly *Erwinia herbicola*) (Aydin et al., 1985). These NRPs were first identified in 1980 as antifungal acyl peptide antibiotics produced by *E. herbicola* (Winkelmann et al., 1980). Their structure consists of eight amino acids which create a cyclic peptide connected to a fatty acid chain, a dehydrobutyric acid, and a sugar molecule (only in herbicolin A). Herbicolin B is a deglycosylated intermediate formed during biosynthesis of herbicolin A. These lipopeptide antimicrobials are active against a broad spectrum of pathogenic fungi and the biosynthetic gene cluster for herbicolin A was identified recently (Matilla et al., 2023; Xu et al., 2022). Herbicolin A acts by interacting with ergosterol, thus disturbing the integrity and permeability of ergosterol-rich fungal cell membranes (Bukrinsky et al., 2020; Xu et al., 2022).

NRP biosynthetic gene clusters encode enzymes that direct the assembly of amino acids and other connected monomers into peptide-like structures (Calcott and Ackerley, 2015; Miyanaga et al., 2022). Specifically, they are composed of the condensation domains, which catalyze the formation of peptide bonds in non-ribosomal peptide biosynthesis, the adenylation domains which are responsible for selective amino acid recognition and activation, and the thiolation domains, which are the carrier proteins. Thiolation domains are responsible for the posttranslational activation of adenylation domains, and their substrates can experience several modifications, including oxidation, epimerization, methylation, and halogenation. Biosynthesis starts with the post-translational modification of the T domain by reaction with a 4’-phosphopantetheine cofactor. A phosphopantetheine cofactor acts as a ‘swinging arm’ for the addition of activated amino-acid and fatty acid groups. The adenylation domain then recognizes and activates the amino acids and the condensation domain catalyzes consequent peptide bond formation. Before interacting with the condensation subunits, the thiolation domains perform modifications, such as epimerization, to the substrate. Finally, the interaction of the peptide chain with a thioesterase domain contributes to the release of the product through hydrolysis or cyclization of the molecule.

According to an old Irish tradition, hanging “Cuileann” (plants of the species *Ilex aquifolium* commonly known as “holly”) inside stables protects animals from getting infected by fungi (ringworms). Although some natural sources of antifungal compounds, such as such as surfactins, iturins, fengycins, syringomycins, hassallidins (Fewer et al., 2021; Meena and Kanwar, 2015), have already been described, current research is still limited regarding the identification and functional characterization of microbially derived antifungal agents that have been shown to be effective, stable, and safe for use in food applications. Following a screening of 610 isolates from “holly” leaves and berries, we identified *P. agglomerans* APC 4211, a strain with strong antifungal activity against food spoilage fungi. By combining genomic and MALDI-TOF MS analysis we revealed that the antifungal activity was due to putative herbicolins A and B. Optimization of antifungal activity was achieved by testing 22 different fermentation media supplemented with different carbon and nitrogen sources. Purified putative herbicolin A has potent activity that may be suitable for industrial applications, including MIC values ranging from 0.2 to 1.5 μg/mL for some target fungal and yeast spoilage organisms, and is also thermally stable and resistant to proteinases. Moreover, we provide evidence of the safety of the molecule against S9 lung epithelial cell lines at concentrations of 50 μg/mL. Furthermore, the stability and effectiveness of the anti-fungal activity of putative herbicolin A was confirmed using skim milk as a model of dairy foods. Together, our findings address existing gaps in the identification and validation of natural antifungal agents and demonstrate that herbicolin A is a promising biocontrol compound with potential use as a safe, clean-label biopreservative in food systems.

## 2. Methods

### 2.1 Bacterial strain selection and culture conditions

*P. agglomerans* APC 4211 (KH-2882) was isolated from a male tree of the native Irish variety of *I. aquifolium* from Bandon Town in County Cork, Ireland. The screening procedure involved homogenization of leaves (5 grams) in 45 ml of sterile Maximum recovery diluent (MRD, Merck™, Germany) using a Stomacher® 400 Circulator Lab Blender (West Sussex, UK) at 300 rpm for 5 minutes. Homogenized samples were serially diluted (1:10 factor) in MRD. Streaking and spreading (100 μl) methods were applied to each dilution in plates with the following media: **a)** tryptic soy broth (TSB, Merck™, Germany); **b)** BD Difco™ Lactobacillus MRS (MRS, Thermo Fisher Scientific, Denmark); and **c)** M17 broth (Merck™, Germany) supplemented with: **i)** lactose (Merck™, Germany) (LM17); **ii)** sucrose (Merck™, Germany) (SM17), **iii)** glucose (Merck™, Germany) (GM17); **iv)** fructose (FM17, Merck™, Germany). The selection of the strain of interest was performed following a screening of 610 isolated bacterial strains against the following fungal indicators: *Candida parapsilosis* UCC, *Saccharomyces cerevisiae* type strain Sa-07140, *Zygosaccharomyces bailii* strain Sa-1403, *Yarrowia lipolytica* strain 78-003, *Fusarium solani* DSM 10696, *Fusarium solani* DSM 1164, *Acremonium persicinum* DSM 1052, *Alternaria alternata* DSM 1102, *Scopulariopsis brevicaulis* DSM 1218, *Geotrichum candidum* KH-1730, *Aspergilus niger* UCC, *Byssochlamys nivea* UCC, *Paecilomyces variotii* UCC, *Cladosporium sp*. UCC, *Phoma sp*. UCC, and *Penicillium notatum* UCC, using the spot-on lawn technique (Zou et al., 2018), as described previously (Kamilari et al., 2025). The growth conditions of each fungal indicator used in this assay are shown in **Table 1**. The growth media used include Sabouraud Dextrose Agar (SDA, Merck, Darmstadt, Germany), Potato Dextrose Agar (PDA, Merck, Darmstadt, Germany), and V8 fruit juice agar (Himedia, Maharashtra, India). Antifungal activity was evaluated based on the diameter of the inhibition zone and was classified into five categories: a) no activity (0 mm); b) weak activity (0.5 – 9 mm); c) moderate activity (10 – 15 mm); d) good activity (16 – 20 mm); e) strong activity (>20 mm). All isolates were tested in triplicate. *P. agglomerans* APC 4211 was cultivated in M17 broth (Merck™, Germany) supplemented with 0.5% sucrose (Merck™, Germany) (SM17) (1% inoculum), and incubated aerobically at 30 °C, under shaking at 200 rpm.

**Table 1.** Growth conditions of selected fungal indicators.

| Fungal indicator | Growth conditions |  |  |
| --- | --- | --- | --- |
|  | Temperature | Time | Medium |
| *Acremonium persicinum* DSM 1052 | 30 °C, aerobic | 5 days | PDA |
| *Alternaria alternata* DSM 1102 | 30 °C, aerobic | 5 days | V8 fruit juice agar |
| *Aspergillus niger* UCC * | 30 °C, aerobic | 3 days | SDA |
| *Byssoschlamys nivea* UCC * | 30 °C, aerobic | 5 days | SDA |
| *Candida parapsilosis* UCC * | 30 °C, aerobic | 24 hours | SDA |
| *Cladosporium* sp. UCC * | 30 °C, aerobic | 5 days | SDA |
| *Fusarium solani* DSM 10696 | 30 °C, aerobic | 5 days | SDA |
| *Fusarium solani* DSM 1164 | 30 °C, aerobic | 5 days | SDA |
| *Geotrichum candidum* KH-1730 | 30 °C, aerobic | 24 hours | SDA |
| *Geotrichum* sp. UCC * | 30 °C, aerobic | 3 days | SDA |
| *Penicillium notatum* UCC * | 25 °C, aerobic | 5 days | SDA |
| *Phoma* sp. UCC * | 25 °C, aerobic | 5 days | SDA |
| *Saccharomyces cerevisiae* type strain Sa-07140 | 30 °C, aerobic | 24 hours | SDA |
| *Scopulariopsis brevicaulis* DSM 1218 | 30 °C, aerobic | 5 days | V8 fruit juice agar |
| *Yarrowia lypolitica* strain 78-003 | 30 °C, aerobic | 24 hours | SDA |
| *Zygosaccharomyces bailii* strain Sa-1403 | 30 °C, aerobic | 24 hours | SDA |

\* Indicators were sourced from APC Microbiome, University College Cork. SDA: Sabouraud Dextrose Agar (Merck, Darmstadt, Germany). PDA: Potato Dextrose Agar (Merck, Darmstadt, Germany). V8 fruit juice agar (Himedia, Maharashtra, India).

### 2.2 Cell-free supernatant (CFS) anti-fungal activity assay

*P. agglomerans* APC 4211 was inoculated in 1L of SM17 medium (1% inoculum) and incubated aerobically in a conical flask at 30 °C, shaking at 200 rpm for 48 hours. A 10 ml sample from the growing culture was collected every 4 hours and centrifuged at 8,000×g at 4°C for 20 min to separate the cells from the supernatant. The supernatant was filtered by passing through a 0.2 μm sterile filter (Sarstedt AG & Co, Numbrecht, Germany) and the CFS was collected. The cell pellet acquired through centrifugation was mixed with 20 ml 70% (v/v) propan-2-ol-containing 0.1% (v/v) trifluoroacetic acid (TFA) (IPA) and incubated at 30 °C for 3 hours under shaking conditions. Following centrifugation at 4,500×g at 4°C for 10 min, the cell pellet IPA-treated CFS (CP) was collected and filtered as described above. To assess the anti-fungal activity of the CFS and the CP the well diffusion assay (WDA) was used as described previously (Kamilari et al., 2025). In summary, a 50 ml 0.75% w/v “sloppy” agar was inoculated with 10^5^ to 10^6^ spores/ml of fungi and placed in a square petri dish. Using a sterilized glass Pasteur pipette (VWR International Limited, cat no. 612-3813), wells of 6 mm diameter were created in the agar plate. The wells were filled with 50 μl of CFS. Following incubation according to the appropriate temperature for each indicator (**Table 1**) the antifungal activity was evaluated based on the diameter of the inhibition zone, as described above.

### 2.3 Whole genome sequencing

Genomic DNA was extracted using the DNeasy® PowerFood® Microbial Kit (MoBio Laboratories Inc., Carlsbad, CA, United States), following the manufacturer’s instructions. The genome of *P. agglomerans* APC 4211 was sequenced by MicrobesNG (Birmingham, UK), using MiSeq Illumina and Oxford Nanopore sequencing platform. Genome assembly, annotation, and prediction of open reading frames (ORFs), ribosomal RNA genes, and transfer RNA genes, were performed as described previously (Kamilari et al., 2025). Additionally, genome assembly was performed using Unicycler version 0.4.3 open-source software (Wick et al., 2017) and visualized using Bandage (Wick et al., 2015). The identification of bacterial plasmids was performed using the Platon software (Schwengers et al., 2020) and visualized using Proksee (Grant et al., 2023). Genome average nucleotide identity (ANI) calculations were performed with pyani (https://github.com/widdowquinn/pyani). The BAGEL4 software (Van Heel et al., 2018) and the antiSMASH v.7.0 software (Blin et al., 2021) were used for bacteriocin/antimicrobial/secondary metabolites gene cluster prediction. The visualization of the non-ribosomally produced peptides (NRP) gene cluster was performed using CAGECAT (https://cagecat.bioinformatics.nl/). The Galaxy platform abricate v. 1.0.1 was used to identify the presence of antibiotic resistance genes and antimicrobial and virulence genes (Seeman, 2020). Functional analysis and domain prediction of the gene cluster encoding for herbicolin A genes from *P. agglomerans* APC 4211, *P. agglomerans* ZJU23, and *P. agglomerans* 9Rz4 were performed using InterPro (https://www.ebi.ac.uk/interpro/). The genome sequence was deposited in GenBank under the accession number SUB15169161.

### 2.4 Colony MALDI-TOF mass spectrometry

*P. agglomerans* APC 4211 cell extract was assessed for the expression of antimicrobial peptides by correlating the presence of peptide molecular masses with the molecular mass of identified antimicrobials. A loop full of *P. agglomerans* APC 4211 colonies grown on SM17 agar was diluted in 50 µl of 75% (v/v) IPA. Following centrifugation at 6000 g for 10 minutes the IPA extract was retained for MALDI-TOF (matrix-assisted laser desorption/ionization coupled to time-of-flight) mass spectrometry analysis by an iDplus Performance MALDI-TOF mass spectrometer (Shimadzu, Duisburg, Germany), as described previously (Kamilari et al., 2025).

### 2.5 Optimization of the inhibitory activity

*P. agglomerans* APC 4211 was inoculated in 50 ml flasks of 22 different media, 20 of which included M17 broth supplemented with 3% of different carbon sources (glucose (Merck™, Germany), sucrose, lactose (Merck™, Germany), fructose (Merck™, Germany) and glycerol (Merck™, Germany)) and 1.5% different nitrogen sources (tryptone (Merck™, Germany) and yeast extract (Fisher Scientific, UK)), half of them contained extra 0.1% casamino acids (Merck™, Germany), and all contained extra 0.2% magnesium sulfate heptahydrate (Fisher Scientific, UK), and 1% sodium chloride (Merck™, Germany). SM17 medium and tryptic soy broth (TSB, Merck™, Germany), were also tested. Samples were collected at 12, 18, 24, 36, 42, and 48 hours. Results were evaluated using the WDA for both the CFS and the CP. Antifungal activity was evaluated as described above. The time point with the highest inhibition zone for each indicator was recorded. The strains *B. nivea* UCC, *Cladosporium* sp. UCC, and *P. variotii* UCC, were used as indicators to detect the highest inhibition activity in the CFS.

### 2.6 Purification and identification of the different antimicrobial compounds

*P. agglomerans* APC 4211 was inoculated in 50 ml LM17T broth (M17 supplemented with 3% lactose, 1.5% tryptone, 0.1% magnesium sulfate heptahydrate, and 1% sodium chloride) and incubated aerobically in a 200 ml conical flask at 30 °C, for 48 hours, under shaking at 200 rpm. A sterile cotton swap was dampened into the culture, and the culture was spread across the surface of a square petri dish containing SM17 agar until the surface was covered completely with the culture. The process was repeated for 20 square petri dishes. The plates were incubated aerobically at 30 °C, for 48 hours. Following incubation, the plates were placed in the -80° C freezer for 48 hours. After the plates were thawed, the cells were collected and following centrifugation at 10,000×g at 4°C for 20 min, the supernatant was separated from the cell pellet. This technique is called the freeze and squeeze method. The supernatant was passed through a 0.2 μm sterile filter (Sarstedt AG & Co, Numbrecht, Germany) to remove the presence of bacterial cells and the CFS was collected. The cell pellet was treated with 50 ml 75% (v/v) IPA and stirred until suspended at room temperature. Following shaking - for 4 hours, and centrifugation at 4,500×g at 4°C for 10min, the IPA-treated cell extract was filtered, sterilized, and collected. The organic solvents were removed using a rotary evaporator (Buchi Labortechnik AG, Flawil, Switzerland).

Both the CFS and the CP were further purified by passing through a 10g (60ml) Strata C18-E SPE column (Phenomenex, Cheshire, UK) pre-equilibrated with methanol and water. Following washing with 100 ml of 40% ethanol, the peptides were eluted using 50 ml of different concentrations of IPA: a) 10%; b) 20%; c) 30%; d) 40%; e) 50%; and f) 75% (v/v). The IPA elute concentration with the greatest antifungal activity was evaluated using the WDA.

Using a rotatory evaporator, the organic solvent was removed. Both the CFS samples and the CP samples were applied to a semi-prep Jupiter C18 reversed-phase HPLC column (10 x 250 mm, 5µ, 300 Å) running a 35-75% gradient over 40 minutes where the mobile phase A was water containing 0.1% TFA, and the mobile phase B was 67% propan-2-ol / 33% acetonitrile containing 0.1% TFA. Eluent was monitored at 214 nm and fractions were collected at 1-minute intervals. The active fractions were analyzed for the antimicrobial mass of interest using a MALDI-TOF mass spectrometer (Shimadzu Europa GmbH, Duisberg, Germany).

### 2.7 Evaluation of the effect of proteases and the stability at different temperatures of the antifungal activity of P. agglomerans APC 4211

The sensitivity of the antifungal CFS of *P. agglomerans* APC 4211 to proteolytic degradation was estimated using proteinase K (1 mg/ml, Sigma-Aldrich, St Louis, USA) and trypsin (1 mg/ml, Merck KgaA, Darmstadt, DE). Specifically, 200 μl of CFS was mixed with **a)** 100 μl of water, and **b)** 100 μl of proteinase K (20 μg/mL), or trypsin (1 μg/mL). Following incubation for 3 hours at 37 °C, each mixture was separated into two eppendorf tubes (150 μl each) and one of the two tubes was incubated for 10 min at 100°C. This causes the inactivation of the proteolytic enzymes. The heat stability of the antifungal CFS of *P. agglomerans* APC 4211 was estimated after incubation for 15 minutes at different temperatures (50 °C, 60 °C, 70 °C, 80 °C, 90 °C, 100 °C, and 121 °C). The activity of the CFS after treatments were tested against *C. parapsilosis* and *Byssochlamys nivea* UCC using the well diffusion assay.

### 2.8 Estimation of the MIC of purified herbicolins against fungal indicators

The MIC of purified herbicolins against yeast and molds was tested in 96-well microtiter plates (Sarstedt, Co. Wexford, Ireland) using a Stratus Microplate Reader (Cerillo, Virginia, USA) by measuring the optical density at 600 nm (O.D._600_). To evaluate fungal spore concentration/ml, the spores were diluted in trypan blue stain (0.4%), which allows their visualization and separation of live from dead cells. Their enumeration was performed using a Hemocytometer (Marienfeld Superior, Lauda-Königshofen, Germany). At each side of the hemacytometer, 10 µl of spore solution was added. The enumeration of spores was performed under a microscope (VWR® VisiScope® Digital Microscope, Dublin, Ireland). To identify sample deposition, the lower magnification (10X or 20X) objective lens was used. Spore counting was performed at a minimum magnification of X400. The calculation of the spore concentration was performed using the following formula:

Total cells/ml = (Total cells counted x Dilution factor x 10,000 cells/ml)/ Number of squares counted

Purified herbicolin was diluted in 1 ml water to create 10 μM stock solution. To a 96-well plate, 100 μl of each fungi growing broth medium was added to each well and 100 μl of 10 μM/ml herbicolin was added to the top row of well wells (A1 to A9). Then, a two-fold serial dilution was performed vertically down the plate starting from the first well, until the 8^th^ well (A to H). Finally, 100 μl of grown culture containing 1 x 10^5^ CFU/ml of yeast cells or 1 x 10^5^ spores/ml was added to all wells. Optical density at 600 nm was measured after 24 hours to 5 days, depending on the fungal indicator growth rate. The MIC was evaluated as the minimum concentration that would prevent the indicator strain from growing. The MIC for each fungal indicator was tested in triplicate.

### 2.9 Comparison of the MIC of purified herbicolin A, purified herbicolin B and combination of purified herbicolins A and B against fungal indicators

The MICs of purified herbicolin A, purified herbicolin B and a combination of herbicolins A and B (as expressed by the strain) were evaluated against the fungal indicators *A. niger* UCC, *B. nivea* UCC, *Geotrichum sp. UCC*, and *Penicillium notatum* UCC as described above.

### 2.10 Comparison of the MIC of purified herbicolins with the MIC of the antibiotics natamycin, fluconazole, itraconazole, ketoconazole, and terbinafine against fungal indicators

Terbinafine hydrochloride in powdered form was dissolved in absolute ethanol to create a stock solution of 10 mg/mL. Terbinafine hydrochloride stock solutions and stock solutions of the antifungal agents fluconazole, itraconazole, and ketoconazole at a concentration of 2 mg/mL were diluted in distilled water, to create working dilutions of 40 μg/ml. Natamycin in powdered form was dissolved in sterile distilled water to create a stock solution of 50 mg/mL The MIC against the fungal indicators *C. parapsilosis* UCC, *Z. bailii* Sa-1403, *A. persicinum* DSM 1052, *A. alternata* DSM 1102, *S. brevicaulis* DSM 1218, *Geotrichum sp. UCC, A. niger* UCC, and *P. notatum* UCC were evaluated as described above. Visualization of the results was performed using a ggplot2 package (Wickham, 2009) v2.2.1 in RStudio (Team, 2015).

### 2.11 Evaluation of the anti-fungal activity of the chemicals sorbic acid, propionic acid and 4-aminobenzoic acid against fungal indicators

The chemicals sorbic acid, propionic acid, and 4-aminobenzoic acid in powdered form were dissolved in water (initially in 1 ml absolute ethanol) to create a working solution of 6000 ppm (6 mg/mL). The MIC against the fungal indicators *Z. bailii* strain Sa-1403, *A. niger* UCC, *Geotrichum* sp UCC, and *P. notatum* UCC was evaluated as described above. Each experiment was performed in triplicate.

### 2.12 S9 cell viability assay

Human airway epithelial cells (S9) were obtained from the American Type Culture Collection (LGC Promochem, Wesel, Germany) (no. CRL-2778). To ensure S9 cells attachment to T75 flasks (Sarstedt, Germany), flasks were treated with 10 ml of “coating solution” which contained fibronectin (1mg/mL, Sigma-Aldrich, St Louis, USA), 0.1% Collagen (30ug/mL, Sigma-Aldrich, St Louis, USA), 35% bovine serum albumin (BSA, 0.01ug/mL, Sigma-Aldrich, St Louis, USA) and LHC-8 medium (Thermo Fisher Scientific, Denmark), and allowed to air dry for 2-3 hours. Cells were cultured in T75 flasks which contained 48 ml of LHC-8 Medium (Thermo Fisher Scientific, Denmark) supplemented with 10% fetal bovine serum (FBS) (Sigma-Aldrich, St Louis, USA), and 1% penicillin (100 units/mL, Sigma–Aldrich, St Louis, USA), and streptomycin (0.1 mg/mL, Sigma–Aldrich, St Louis, USA) at 37 °C/5% CO_2_. Feeding with fresh medium was performed every two days, and cells were checked under a microscope until they reached about 80% of cell viability. To measure cell viability, cells were washed with 10 ml of phosphate-buffered saline (PBS) (Sigma–Aldrich, St Louis, USA) and detached from the flask using 2mL trypsin (Sigma– Aldrich, St Louis, USA) and incubated at 37 °C for about 7 mins. Then, 50 μL of cells in trypsin were mixed with 50 μL trypan blue. 10 μL of the mixture was added into a countess (Invitrogen™, Thermo Fisher Scientific, Denmark), and the number of total cells, live cells, dead cells, and % of viability were measured. In a 96-well plate previously treated with “coating solution” 1 × 10^5^ cells/well/200 µL supplemented LHC-8 medium were added. Cells were treated with four different concentrations of herbicolin A and herbicolin B (0.5, 2.5, 10, and 50 μg/ml). A negative control included no treatment, and a positive control included triton X (1%). Five technical replicates were used for each treatment. Results were expressed as cell viability percentage (%) and analysed using a one-way ANOVA to assess differences among the control and herbicolin A and B concentrations. When the ANOVA indicated a significant overall effect, post hoc pairwise comparisons between groups were performed using Student’s *t*-tests with Holm correction for multiple testing. Effect sizes and 95% confidence intervals were calculated. Statistical analyses were performed and visualized using the ggstatsplot package (Patil, 2021) in RStudio (Team, 2015).

### 2.13 Skim milk as dairy food model

The fungal strains *A. niger* UCC, *Penicillium notatum* UCC, and *Geotrichum* sp UCC were tested against three different concentrations of herbicolin A (0.5, 2.5, and 5 μg/ml) using 20 ml of 10% skim milk as a dairy food model. Fungi at a concentration of 10^3^ spores/ml were added to conical flasks containing 20 ml pasteurized milk containing different concentrations of lipopeptides and incubated at 30 °C under shaking at 150 rpm. The inhibitory activity of the lipopeptide against fungi was determined over time following 24 hours, 48 hours and 72 hours of incubation. Liquid samples were serially diluted (1:10 factor) in MRD and 100 μl of each sample was spread in Sabouraud Dextrose Agar with Chloramphenicol (Sigma-Aldrich, St Louis, USA). The number of fungal cells in CFU was evaluated following 48 hours of incubation at 30 °C. Statistical differences in CFU among different concentrations of herbicolin A for each indicator were analyzed using the ggstatsplot package (Patil, 2021) in RStudio (Team, 2015) as described above. Each experiment was performed in triplicate.

## 3. Results

### 3.1 Antifungal activity of *P. agglomerans* APC 4211

*P. agglomerans* APC 4211 was selected as the strain with the greatest antifungal activity against specific fungal indicators following a screening of 610 strains isolated from “holly” leaves and berries. Using spot-assays, the antifungal activity of *P. agglomerans* APC 4211 against yeast and molds was evaluated. The strain was active against all the tested indicators except *Fusarium solani* DSM 10696. The most sensitive fungus was *Byssochlamys nivea* UCC (**Table 2**). The CFS of *P. agglomerans* APC 4211 was also assessed for antifungal activity against the indicators listed above using the well-diffusion assay. The strongest inhibitory activity was observed against *B. nivea* UCC while good activity was also shown against *P. variotii* UCC and *Cladosporium sp*. UCC. Furthermore, the CFS indicated negligible activity against the yeasts *S. cerevisiae* type strain Sa-07140, *Z. bailii* strain Sa-1403 nor *Y. lipolytica* strain 78-003.

**Table 2.** Antimicrobial activity of *P. agglomerans* APC 4211 against indicator strains using spot assay (using the strain), well diffusion assay (using the CFS), well diffusion assay (using the semi-purified CFS) and MIC.

| Indicator | Inhibition |  | Partially purified peptide inhibition zone (mm) | MIC (µg/ml) |
| --- | --- | --- | --- | --- |
|  | Spot assay | WDA |  |  |
| *Acremonium persicinum* DSM 1052 | ++++ | +++ | 25 mm | 0.3 |
| *Alternaria alternata* DSM 1102 | ++++ | +++ | 35 mm | 1.3 |
| *Aspergillus niger* UCC | ++ | + | 32 mm | 0.9 |
| *Byssoschlamys nivea* UCC | ++++ | ++++ | 41 mm | 1.5 |
| *Candida parapsilosis* UCC | ++++ | ++ | 20 mm | 1.2 |
| *Cladosporium sp.* UCC | +++ | +++ | 35 mm | 1.5 |
| *Fusarium solani* DSM 10696 | - | - | - | - |
| *Fusarium solani* DSM 1164 | ++ | - | 14 mm | 1.3 |
| *Geotrichum candidum* KH-1730 | ++ | ++ | 22 mm | 0.3 |
| *Geotrichum sp.* UCC | ++ | - | 28 mm | 0.4 |
| *Paecilomyces variotii* UCC | +++ | +++ | 39 mm | 0.8 |
| *Penicillium notatum* UCC | ++ | ++ | 35 mm | 0.4 |
| *Phoma* sp. UCC | ++ | ++ | 34 mm | 0.3 |
| *Saccharomyces cerevisiae* type strain Sa-07140 | +++ | - | 28 mm | 0.2 |
| *Scopulariopsis brevicaulis* DSM 1218 | ++++ | +++ | 34 mm | 1.1 |
| *Yarrowia lipolytica* strain 78-003 | +++ | - | 28 mm | 0.2 |
| *Zygosaccharomyces bailii* strain Sa-1403 | +++ | + | 38 mm | 0.2 |

. +: weak activity, inhibition zone 0,5 – 9 mm; ++: moderate activity, inhibition zone 10 – 15 mm; +++: good activity, inhibition zone 16 – 20 mm; ++++: strong activity, inhibition zone >20 mm; -: no activity.

### 3.2 Genomic analysis of *P. agglomerans* APC 4211 reveals a gene cluster encoding herbicolin A

Whole genome sequence analysis *of the P. agglomerans* APC 4211 genome was performed to identify the genetic basis responsible for the observed antifungal activity. The genome assembly was performed using Unicycler version 0.4.3 open-source software resulting in a complete single chromosome of 5,079,259 base pairs and a GC content of 54.7%. Additionally, the analysis detected two plasmids with sizes 129,631 bp (plasmid 1) and 79,899 bp (plasmid 2). These were confirmed using the Platon software (**Figure 1**). Comparison of the whole genome of *P. agglomerans* APC 4211 with 10 other *P. agglomerans* complete circular genomes revealed that the strain exhibits an average nucleotide identity (ANI) ≥98% with all other *P. agglomerans* strains (**Supplementary Figure S1**). The Galaxy platform abricate v. 1.0.1, and the Prokka (Galaxy version 1.14.6) were used to detect the possible presence of virulence factors in the genome (Seeman, 2020; Seemann, 2014). This analysis revealed the absence of virulence factors but confirmed the presence of the gene *oqxB*, the expression of which can be responsible for resistance to nalidixic acid. Additionally, the strain lacked genes that were reported to provide virulence to *P. agglomerans* KM1 strain (Guevarra et al., 2021). These included the absence of the structural gene components of the type VI secretion system (T6SS), such as the valine-glycine repeat G (VgrG) protein. Genes associated with the type III secretion system (T3SS), which encodes for a genotypic trait present in phytopathogenic *P. agglomerans*, were also absent (Rezzonico et al., 2009).

**Figure 1.**
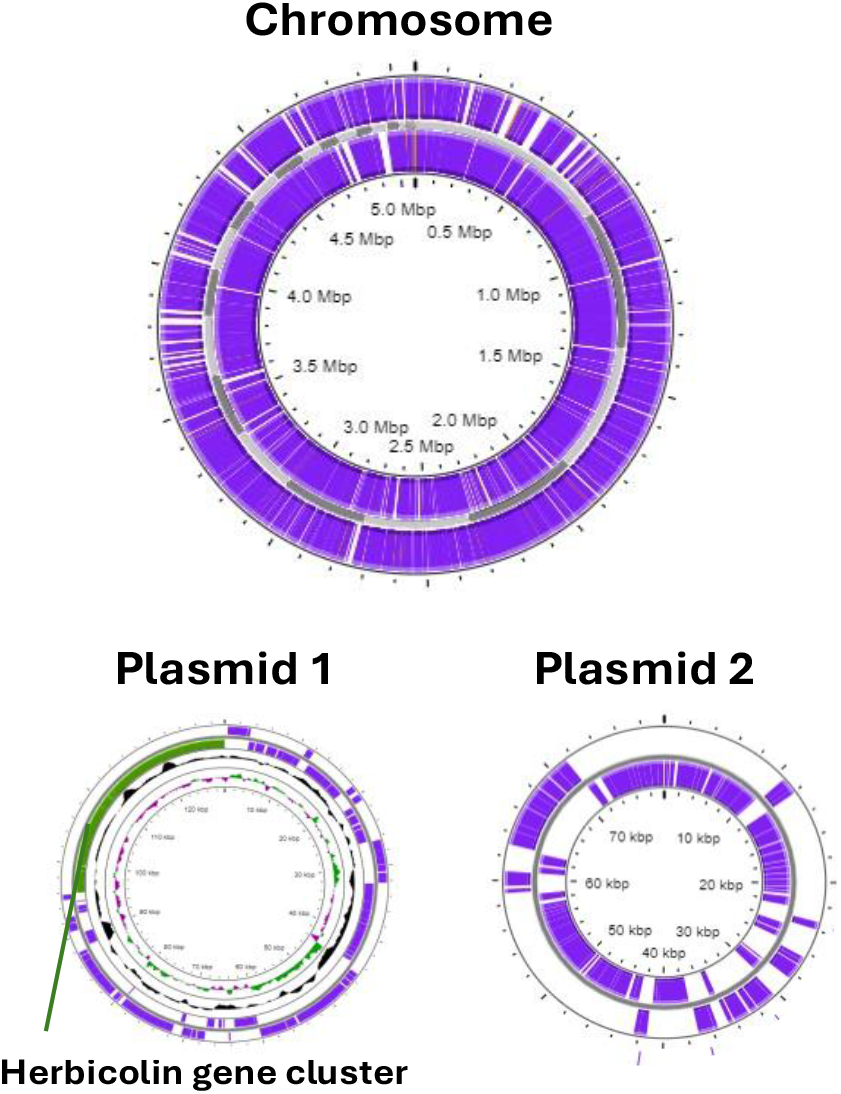
Genomic map of *P. agglomerans* APC 4211 showing the chromosome and the two plasmids. The green section in plasmid 1 indicates the gene cluster encoding for herbicolins A and B. The map was created following genome assembly using Unicycler version 0.4.3 open-source software and visualized using Proksee (https://proksee.ca/).

Genome mining using BAGEL4 and antiSMASH revealed the presence of gene clusters potentially encoding several antimicrobial or bioactive compounds (**Table 3**). Of particular interest was a gene cluster associated with non-ribosomal peptide synthetases (NRPS) with 6% similarity to those encoding for hassallidin C, which is a known antifungal glycolipopeptide (Vestola et al., 2014), according to the antiSMASH software. The gene cluster was bidirectional with a length of ∼42 kb and was identified on plasmid 1 (**Figure 1**). Also, this gene cluster indicated high sequence similarity with a recently published one, which encodes for the lipopeptide herbicolin A (Matilla et al., 2023; Xu et al., 2022). However, the genes encoding HbcA and HbcB as described previously were missing (Matilla et al., 2023; Xu et al., 2022). Exploration for other genes predicted to encode adenylation domain containing proteins located around the putative herbicolin A gene cluster elsewhere on the plasmid in *P. agglomerans* APC 4211 detected only the presence of an adenylation domain predicted to encode for anthranilic acid. High sequence similarity was observed for the structural components herbicolin synthase subunit D (HbcD) (99%), herbicolin synthase subunit E (HbcE) (99%), herbicolin synthase subunit I (HbcI) (99%), an oxidoreductase (HbcF) (100%), an MbtH-like protein (100%), an ABC transporter (100%), a glycosyltransferase (HbcJ) (99%), and a dehydrogenase (98%), previously described by Matilla and coworkers (Matilla et al., 2023) (**Table 4, Supplementary Figure 2**). Furthermore, the gene cluster was compared with the gene clusters encoding for hassallidin C, gramicidin encoded by *Brevibacillus brevis* (Chen et al., 2012), and gramicidin S encoded by *Aneurinibacillus migulanus* (Berditsch et al., 2007), indicating low sequence similarity between herbicolin’s A structural components HbcD, HbcE, and HbcI and the genes hasN, hasO, and hasY respectively, of hassallidin C (**Supplementary Figure 2)**.

**Table 3.**
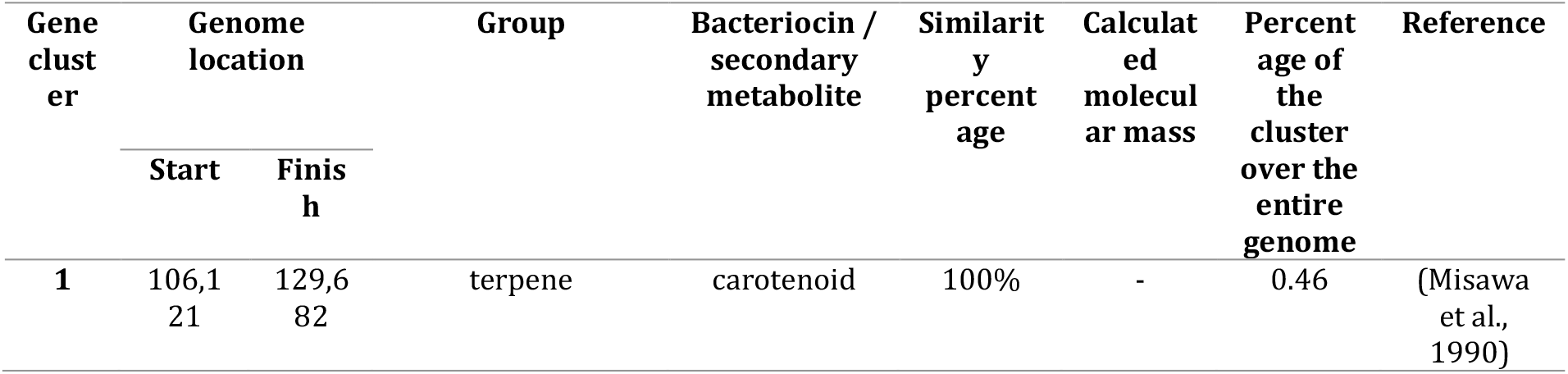
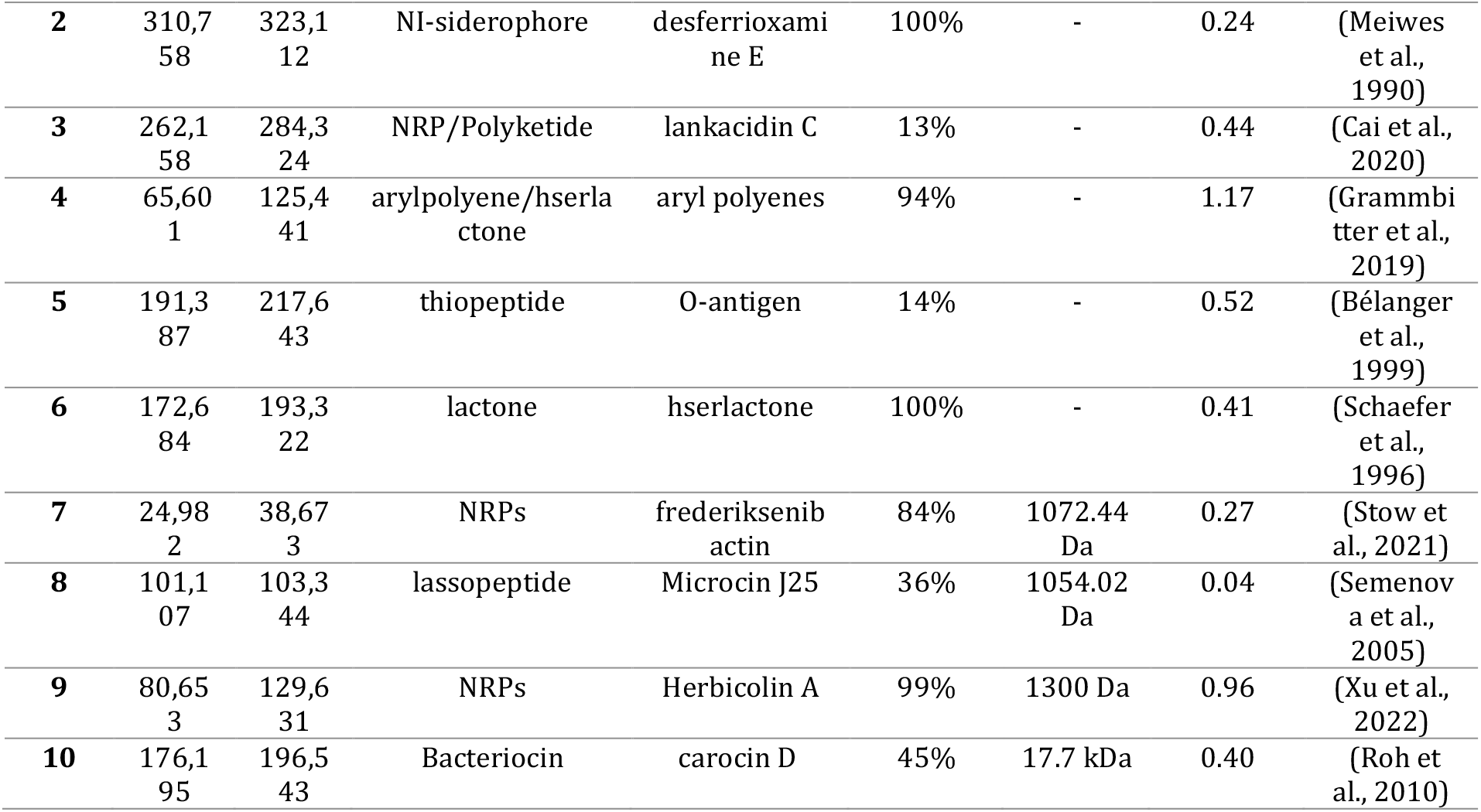
Detected bacteriocins and secondary metabolites in *P. agglomerans* APC 4211 genome using BAGEL4 and antiSMASH.

**Table 4.**
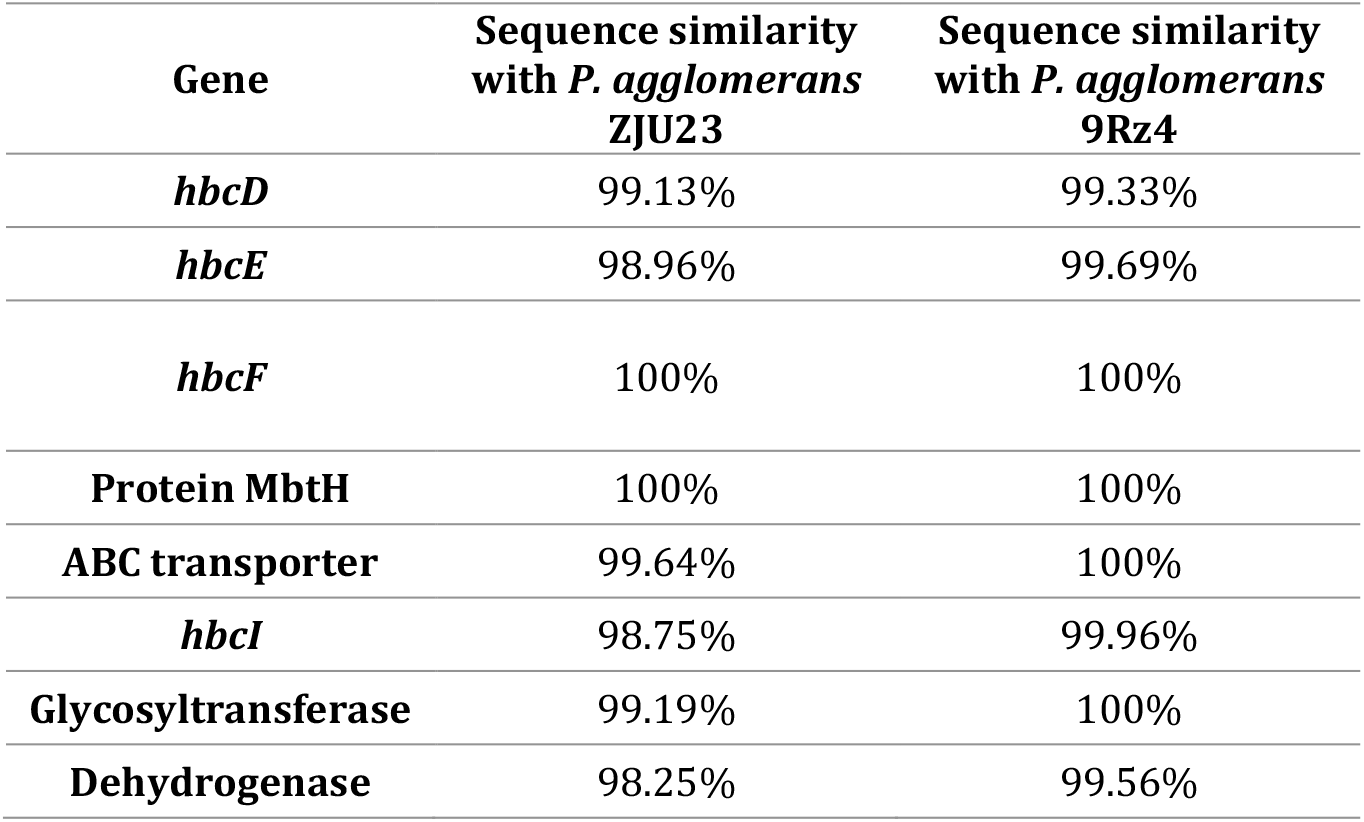
Sequence similarity among the structural components of herbicolin A biosynthetic gene cluster.

**Figure 2.**
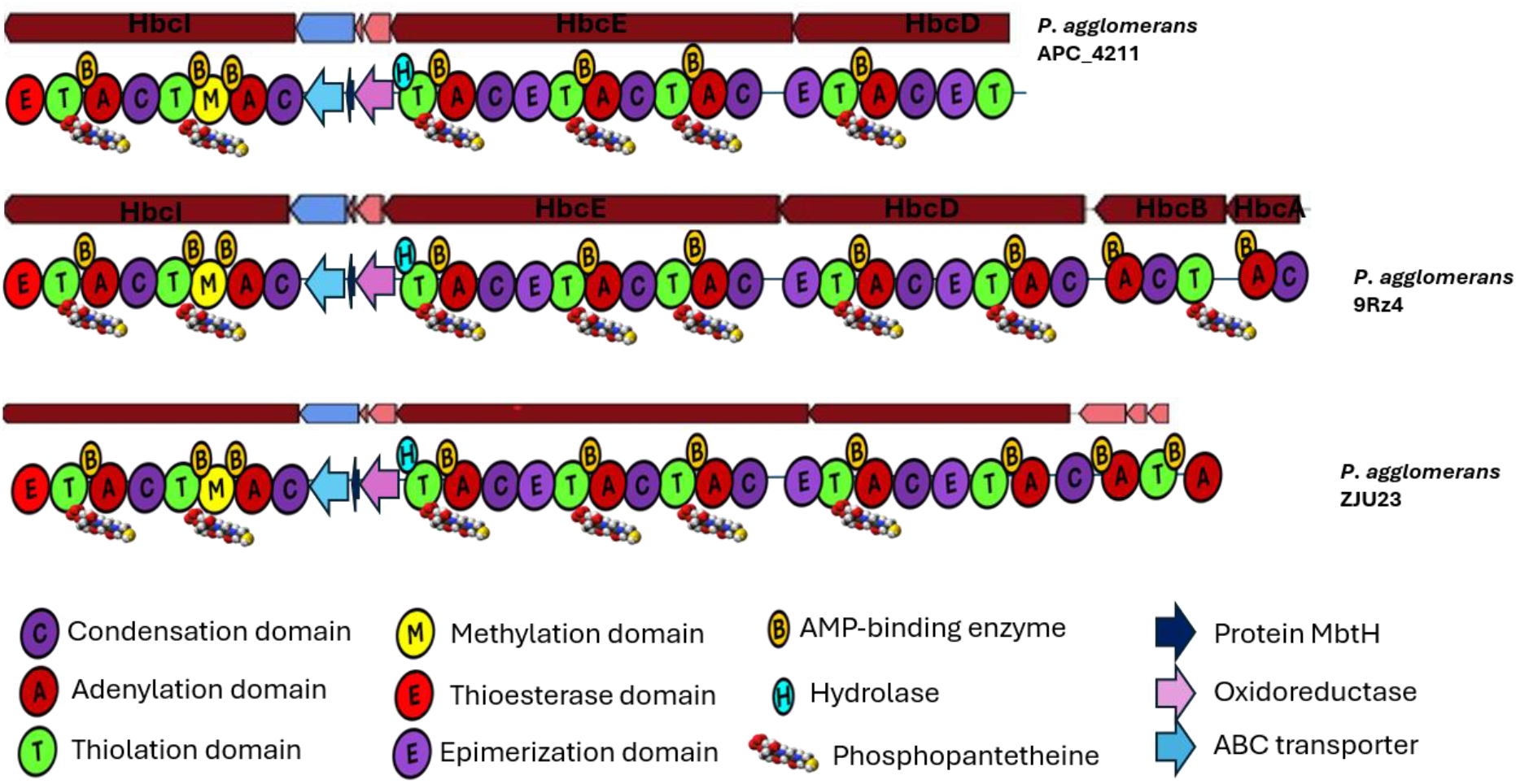
Herbicolin A gene cluster. Domain identification and comparison among the gene clusters associated with herbicolin A from *P. agglomerans* APC 4211, *P. agglomerans* ZJU23, and *P. agglomerans* 9Rz4, using the InterPro database (https://www.ebi.ac.uk/interpro). The gene cluster was predicted to encode Herbicolin A and Herbicolin B.

Protein functional analysis and domain identification using InterPro and antiSMASH, indicated that the protein encoded by the *hbcD* gene from *P. agglomerans* APC 4211 was missing one adenylation domain (A) compared to that encoded by *hbcD* gene from *P. agglomerans* ZJU23 and *P. agglomerans* 9Rz4, and one condensation (C) subunit compared to the protein encoded by the *hbcD* gene from *P. agglomerans* 9Rz4 (**Figure 2)**. In agreement with Matilla and coworkers (Matilla et al., 2023) and Xu and coworkers (Xu et al., 2022) the *hbcE* and *hbcI* genes encoded the same C, A, and T domains. Additionally, the *hbcI* gene encodes a protein containing a putative methyltransferase domain (M), indicating that an amino acid is N-methylated and a thioesterase (E) subunit, which is responsible for the cyclization of the molecule. The *hbcE* gene is followed by a putative gene encoding a 3-oxoacyl-(acyl-carrier-protein) reductase, an enzyme that participates in fatty acid biosynthesis. Following this, a gene encoding for MbtH, a protein detected in NRPs to assist in their folding, stability and function was identified (Zwahlen et al., 2019). Additionally, a gene encoding an ABC transporter ATP-binding/permease protein was found, which is possibly involved in the translocation of molecules from the cell membrane. The *hbcI* gene is followed by a glycosyltransferase which is predicted to catalyze the glycosylation of the molecule. Downstream, a gene predicted to encode for a dehydrogenase is followed by a gene predicted to encode for a multidrug export protein. Following downstream is a gene predicted to encode for a phosphopantetheinyl transferase, an enzyme responsible for the addition of the phosphopantetheine cofactor to the T domain was found. Furthermore, the predicted presence of an *abi* family immunity gene, which possibly encodes self-immunity to the strain, was observed. The presence of a tyrosine-type recombinase/integrase and an Rpn recombination-promoting transposase homologs associated with the gene cluster suggests the possibility of mediating a horizontal transfer of the gene cluster (**Supplementary Figure 3)**.

**Figure 3.**
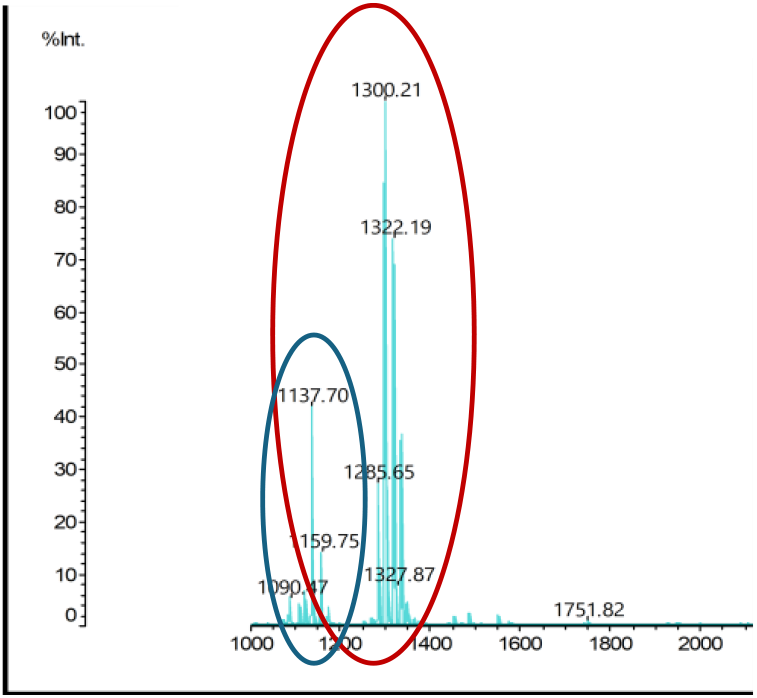
Colony MALDI-TOF mass spectrum showing the detected molecular masses of *m/z* values herbicolin A (1,300 Da), and herbicolin B (1,138 Da).

### 3.3 Colony mass spectrometry

Mass spectra of the cell extract indicated that *P. agglomerans* APC 4211 produced peptides with *m/z* values of 1300.21 Da and 1137.7 Da (**Figure 3**).

### 3.4 Optimization of the inhibitory activity

To optimize the fermentation conditions of *P. agglomerans* APC 4211 with a view to enhancing the antifungal activity in the CFS, we tested 22 different production media supplemented with different carbon sources, nitrogen sources, and the extra addition of casamino acids, magnesium sulfate heptahydrate, and sodium chloride. The greatest inhibitory activity against *B. nivea* UCC, *Cladosporium* sp. UCC and *P. variotii* UCC was observed following incubation for 48 hours in the medium LM17T (M17 supplemented with 3% lactose, and 1,5% tryptone), (inhibition zone diameter 20 mm, 21 mm and 22 mm, respectively) (**Figure 4A, 4B, 4C**).

**Figure 4.**
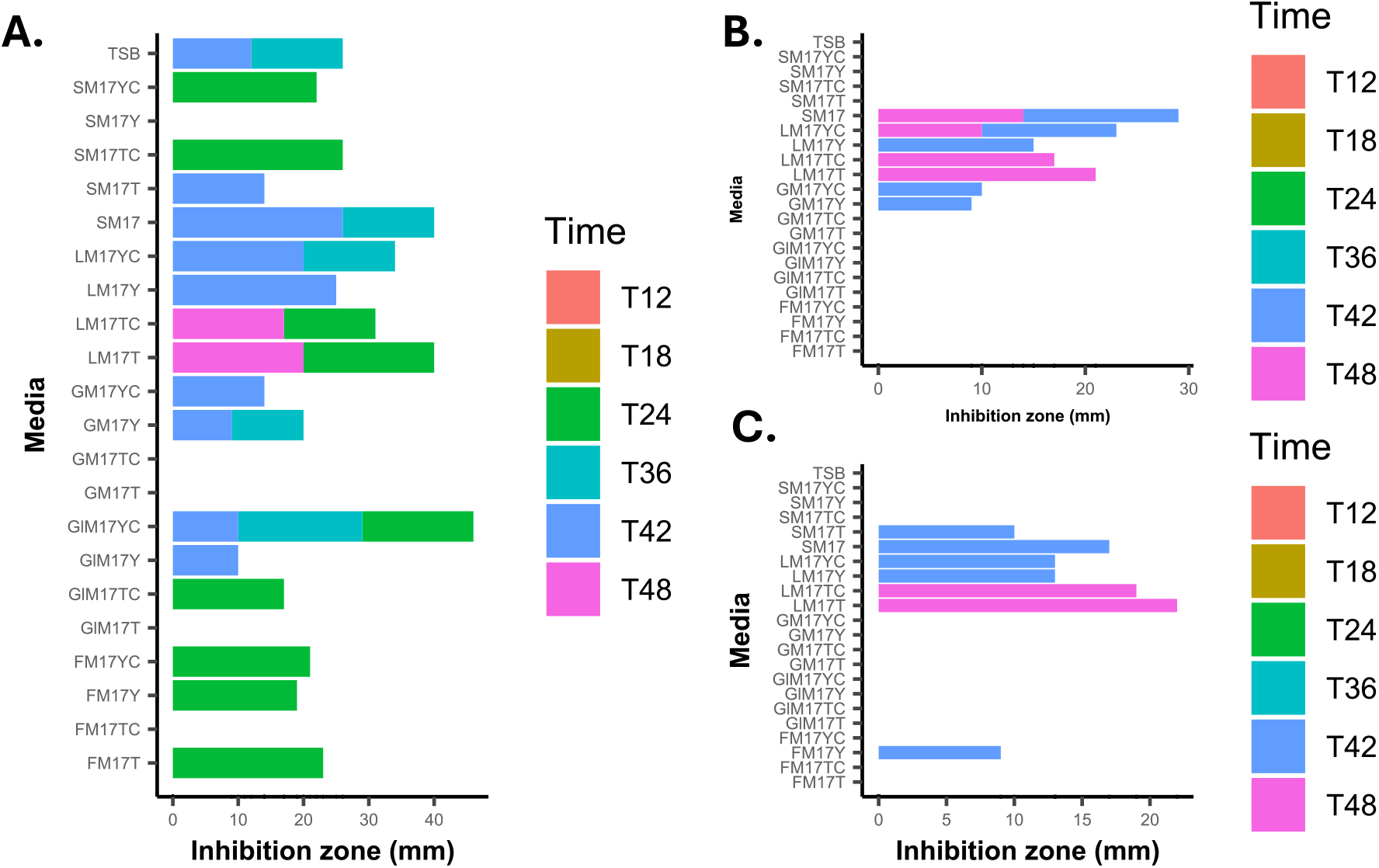
Optimization of *P. agglomerans* APC 4211 inhibitory activity by testing 22 different media against **A.** *B. nivea* UCC; **B**. *Cladosporium* sp. UCC; and **C**. *P. variotii* UCC. Time points 12, 18, 24, 36, 42, and 48 hours are color-coded as follows: T12: coral, T18: yellow/gold, T24: green, T36: turquoise, T42: blue, and T48: pink.

### 3.5 HPLC and MALDI TOF MS analysis identifies an active HPLC fraction containing masses correlating with the theoretical masses of herbicolin A and B

Following C18 column purification, the size of the inhibition zone from the partially purified CFS was larger against all the indicators, except for *F. solani* DSM 10696, the only indicator with no inhibition activity in the spot assay (**Figure 5, Table 2**). The highest activity was observed against *B. nivea* UCC (41 mm). Interestingly, the partially purified CFS was active against the indicators *F. solani* DSM 1164 (14 mm), *S. cerevisiae* type strain Sa-07140 (28 mm), *Y. lipolytica* strain 78-003 (28 mm) and *Geotrichum* sp. UCC (28 mm), for which no inhibition activity had been previously observed using supernatant in the well diffusion assay.

**Figure 5.**
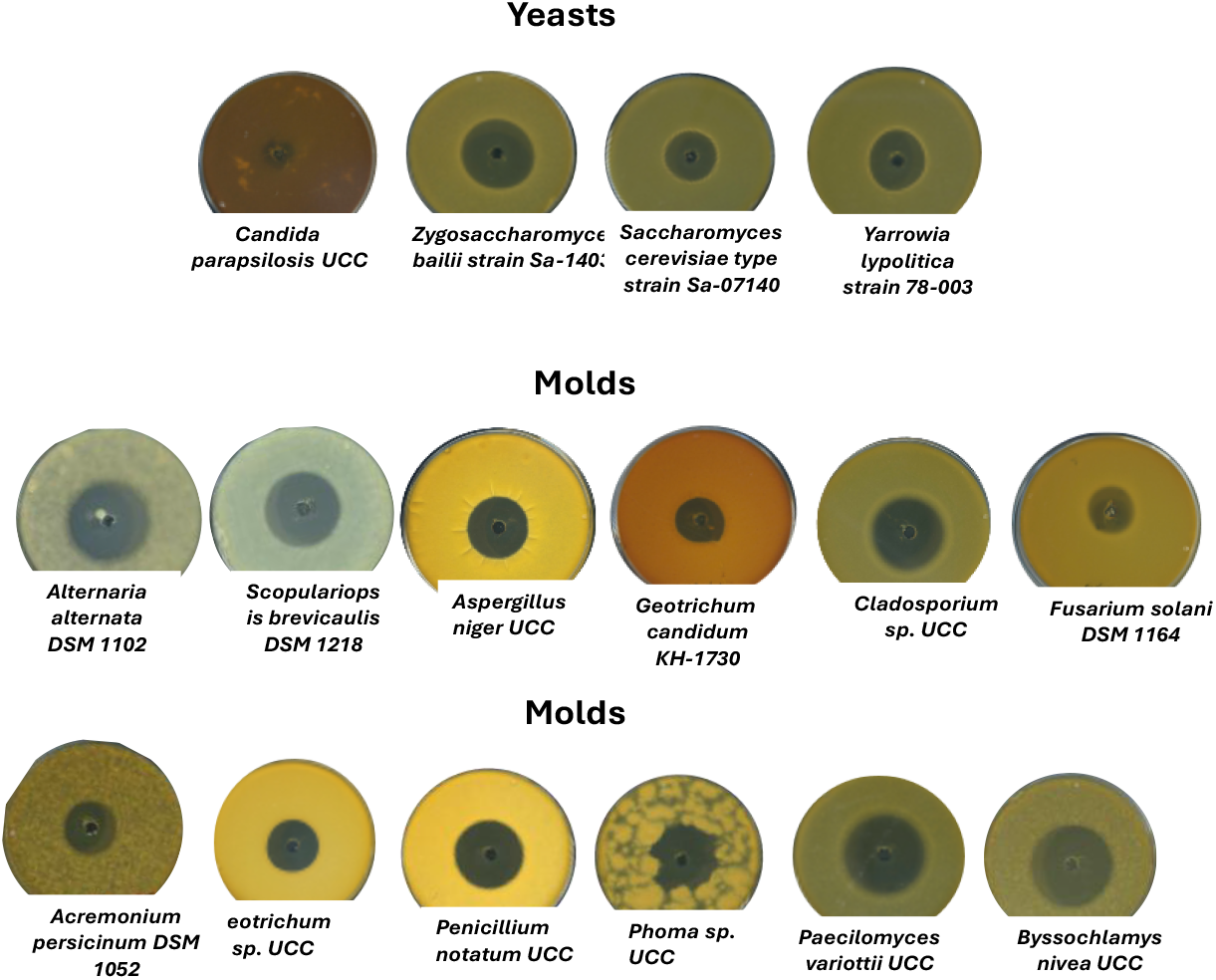
Inhibitory activity of *P. agglomerans* APC 4211 partially purified CFS against mold indicators strains.

The C18 SPE eluents were fractionated by reversed-phase HPLC and assessed for antifungal masses of interest by MALDI TOF MS. The HPLC chromatogram of the partially-purified CFS and the IPA-treated cell pellet indicated elution of a mass correlating with the theoretical masses of the lipopeptide herbicolin A (1300.8 Da) in fractions 30 to 34 (**Figure 6A**). In addition to herbicolin A, a peptide with an additional 18 Da mass (1300.8 Da +18 Da) was detected in fraction 32, indicating that a small percentage of the peptide is modified. Furthermore, in combination with the herbicolin A mass, a mass correlating with the putative herbicolin B mass (1138 Da) was detected in fractions 33 and 34.

**Figure 6.**
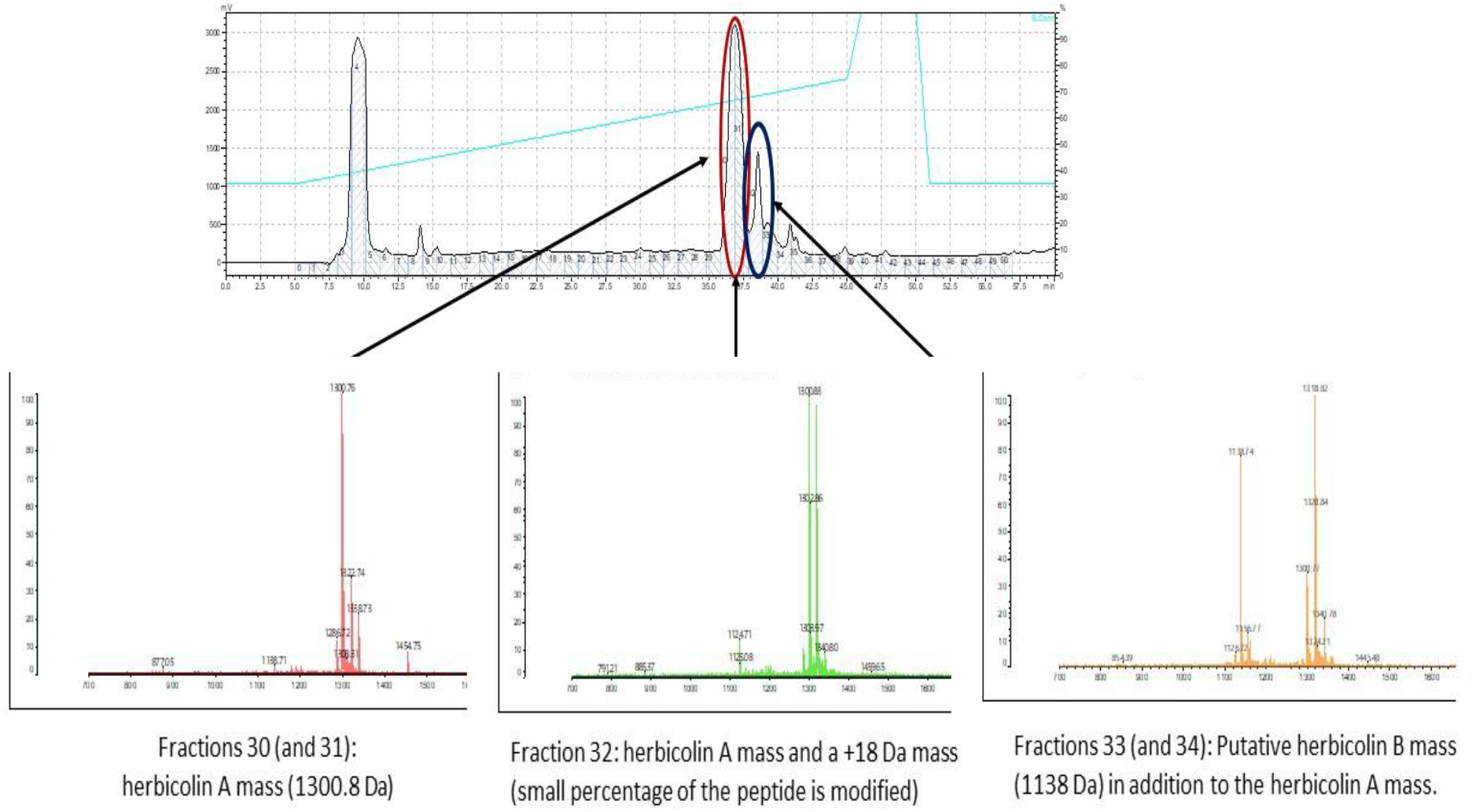
HPLC chromatogram of the CFS of *P. agglomerans* APC 4211 showing elution of herbicolin A in fraction 30, a peptide with an additional 18 Da mass in fraction 32 and herbicolin B in fraction 33.

### 3.6 Stability and Activity of the partially purified CFS to proteases and temperature

The sensitivity of the partially purified CFS to the proteases proteinase K and trypsin degradation and its stability after exposure to different temperatures, ranging from 40 ° C to 100 ° C, for 15 minutes, was estimated using the well diffusion assay. The assay indicated that the antifungal activity of the CFS is stable after exposure to the tested proteases (**Supplementary Figure 4A**) and to the tested temperatures (**Supplementary Figure 4B**).

### 3.7 Estimation of the MIC of purified herbicolins against fungal indicators

The MIC of purified herbicolins A and B was evaluated against the fungal indicators in 24-hours to 5-day experiments (**Table 2**). The duration of the experiments was determined based on the time-point needed for each fungi to grow. The lowest MIC value was detected against the yeasts *S. cerevisiae* type strain Sa-07140, *Y. lypolitica* strain 78-003, and *Z. bailii* strain Sa-1403 (0.2 μg/ml). The MIC value against the other fungi ranged from 0.2 μg/ml to 1.5 μg/ml. No activity was detected against *F. solani* DSM 10696.

### 3.8 Comparison of the activity of herbicolin A with antibiotics natamycin, fluconazole, ketoconazole, itraconazole and terbinafine and chemical preservatives

Initially, the MIC of pooled purified herbicolins A and B (as produced by *P. agglomerans* APC 4211) was compared with the MIC of individual herbicolins A and B. The MIC of pooled herbicolins was lower compared to the MIC of purified herbicolin B, but similar to herbicolin A against the tested fungal indicators (**Supplementary Figure 5**), indicating that herbicolin A has greater antifungal activity compared to herbicolin B.

Herbicolin A was observed to be more effective against *A. persicinum* DSM 1052, *Geotrichum* sp. UCC, and *S. brevicaulis* DSM 1218, with MIC values ranging between 0.08–0.4 μg/ml, compared to the antibiotics fluconazole (5 - >20 μg/ml), ketoconazole (0.4 - >20 μg/ml), itraconazole (0.04 - >20 μg/ml), terbinafine (0.3 - >20 μg/ml), and natamycin (1.125 – 10 μg/ml) (**Figure 7**). Herbicolin A was also observed to inhibit *A. niger* UCC (0.87 μg/ml) and *P. notatum* UCC (0.4 μg/ml) more effectively compared to fluconazole and ketoconazole whose MIC was higher than 20 μg/ml. Additionally, they were equally more effective against Z. *bailii* Sa-1403 than itraconazole, as compared to the other antibiotics.

**Figure 7.**
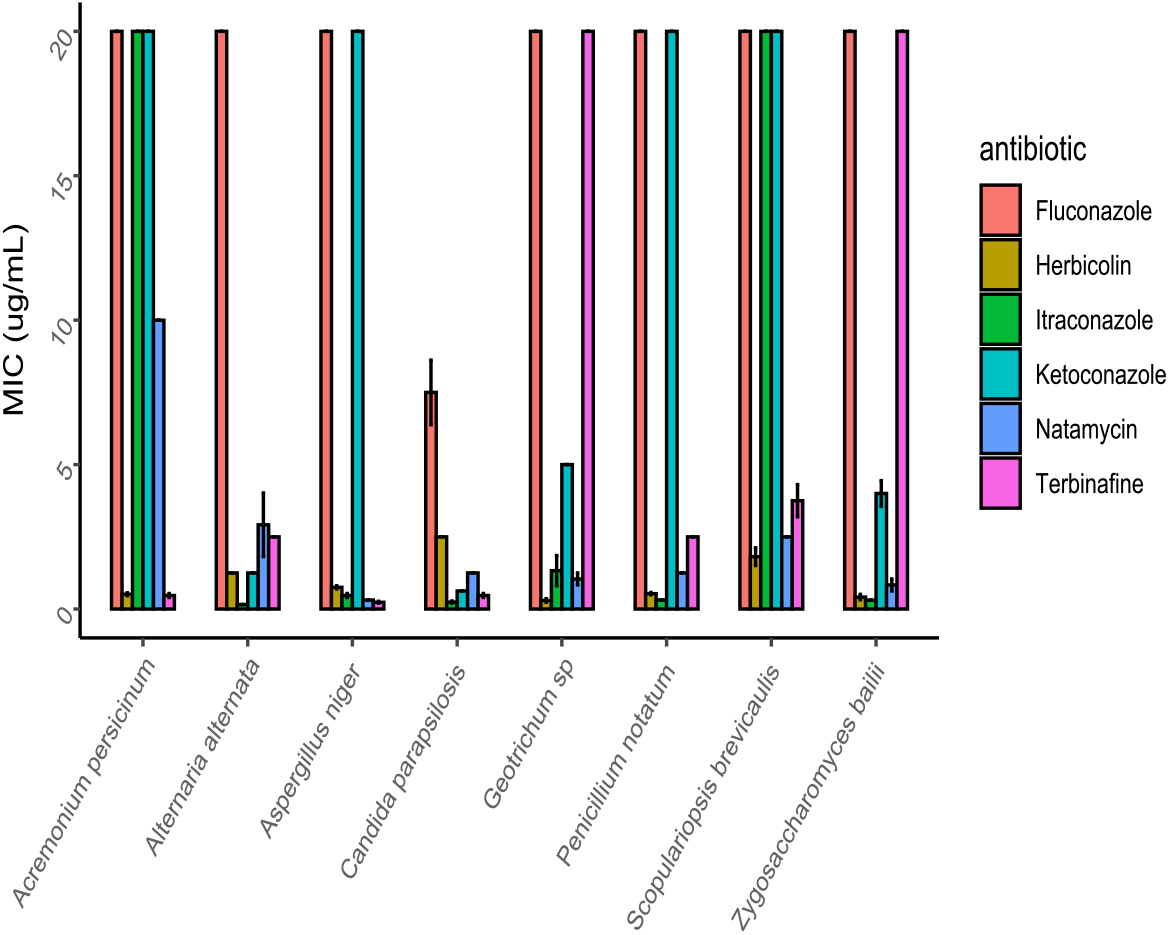
Comparison of fungal indicators MIC values against purified herbicolin A and antibiotics fluconazole, ketoconazole, itraconazole, terbinafine and natamycin.

To evaluate the efficacy of chemical preservatives, the strains *Z. bailii* Sa-1403, *A. niger* UCC, *Geotrichum* sp UCC, and *P. notatum* UCC were tested in the presence of sorbic acid, propionic acid, and 4-aminobenzoic acid. Only sorbic acid indicated inhibitory activity against the tested indicators, in MIC concentrations ranging from 187 ppm (0.187 mg/ml) to 750 ppm (0.75 mg/ml) (**Supplementary Figure 6**). Propionic acid and 4-aminobenzoic acid in a maximum tested concentration of 3000 ppm (3 mg/ml) did not inhibit the growth of the indicators.

### 3.9 Herbicolins A and B are not toxic against S9 human epithelial cells

One-way ANOVA revealed that herbicolins A and B had no toxicity effects on S9 human epithelial cell viability compared to the negative control (no treatment) (**Figure 8**). On the contrary, positive control (triton X) had significant effect on cell viability compared to herbicolins A and B, resulting in cell death following 48 hours of incubation (F(9, 15.63) = 47.49, *p* < 0.001). Post hoc pairwise comparisons using Student’s *t*-tests with Holm correction showed significant differences only between the positive control and the other values, but not between the negative control and the different herbicolins A and B concentrations.

**Figure 8.**
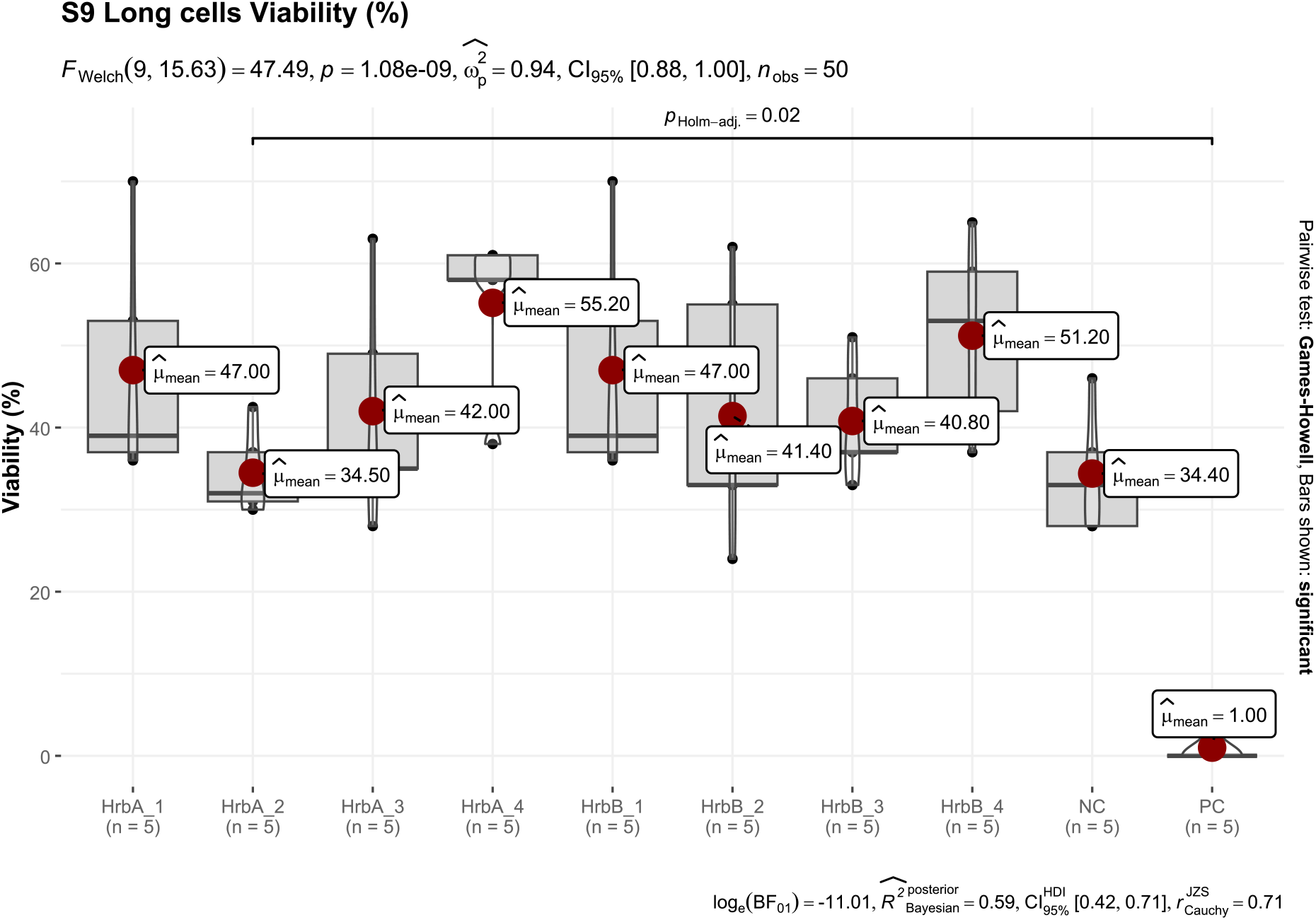
Barplots showing percentage of viability of S9 epithelial cells following administration of different concentrations of herbicolin A (HrbA) and B (HrbB) (1: 0.5, 2: 2.5, 3: 10, and 4: 50 μg/mL). NC: negative control represents no treatment. PC: Positive control represents 1% Triton X. Box-and-whisker plots show the median (horizontal line), interquartile range (box), and minimum and maximum values (whiskers). Data are shown as individual values with group means. Statistical significance was assessed using one-way ANOVA followed by Holm-corrected pairwise Student’s *t*-tests. The equation shown at the top of the plot reveals the statistical model used for comparisons. The values shown at the bottom of the plot indicate the ANOVA results, including the F-statistic with degrees of freedom, the associated p-value, and the effect size with 95% confidence interval.

### 3.10 Skim milk as a dairy food model

To estimate the anti-fungal effects of herbicolins against *A. niger* UCC, *P. notatum* UCC and *Geotrichum* sp UCC (using 10^3^ fungal spores) in a dairy food model, herbicolin A was tested at three different concentrations (0.5, 2.5, and 5 μg/ml). Following 72 hours of incubation with herbicolin A at a concentration of 5 μg/ml no viable cells of *A. niger* UCC and *Penicillium notatum* UCC were recovered, and the viability of *Geotrichum* sp UCC was reduced significantly (2-log_10_ CFU reduction, p=0.04) (**Figure 9**). Administration of herbicolin A at a concentration of 2.5 μg/ml caused a significant reduction in the CFU of *A. niger* UCC, *P. notatum* UCC and *Geotrichum* sp UCC compared to untreated controls (*p* < 0.05). A significant reduction was observed in the CFU of *P. notatum* UCC at a concentration of 0.5 μg/ml (*p* < 0.05), but not against the other fungi.

**Figure 9.**
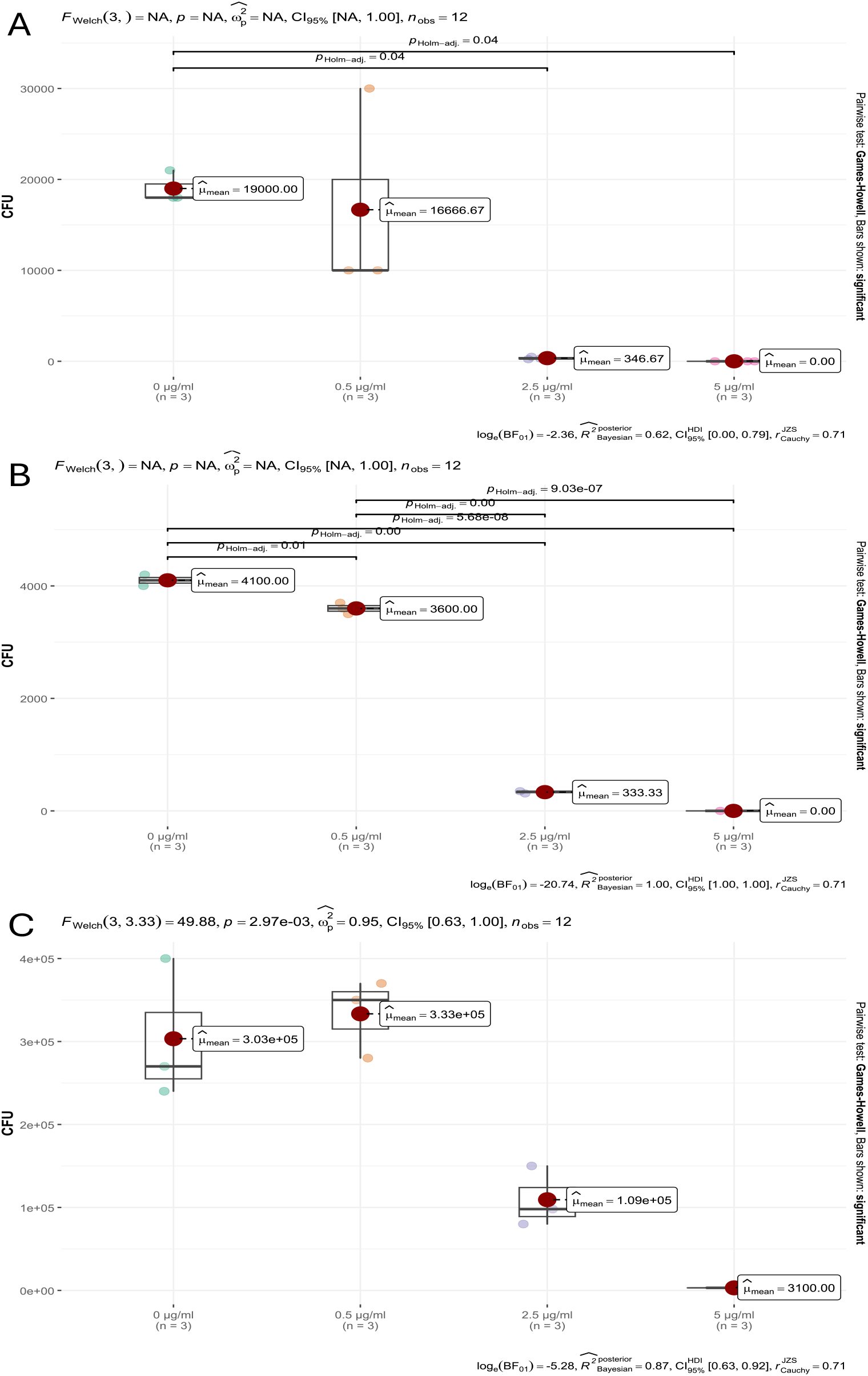
Barplots of measured fungal CFU (initial concentration of 10^3^ spores/mL) per herbicolin A treatment (0, 0.5, 2.5 and 5 μg/mL), following 72 hours incubation in 10% skim milk. CFU per treatment for *A. niger* UCC (A); CFU per treatment for *P. notatum* UCC (B); CFU per treatment for *Geotrichum* sp. UCC (C). Box-and-whisker plots show the median (horizontal line), interquartile range (box), and minimum and maximum values (whiskers). Statistical significance was assessed using one-way ANOVA followed by Holm-corrected pairwise Student’s *t*-tests. The equation shown at the top of the plot reveals the statistical model used for comparisons. The values shown at the bottom of the plot indicate the ANOVA results, including the F-statistic with degrees of freedom, the associated p-value, and the effect size with 95% confidence interval.

## 4. Discussion

The increasing consumer demand for synthetic additives-free and naturally processed foods highlights the importance of the identification of novel, effective, and safe biopreservatives able to prevent the growth of spoilage and pathogenic fungi. AMPs may represent good candidates for an effective solution to this global problem (Martínez-Culebras et al., 2021). In this study, we demonstrated the ability of the strain *P. agglomerans* APC 4211 isolated from “holly” tree leaves to inhibit the growth of spoilage fungi. By applying a combination of whole genome sequencing and genome mining analysis, using antiSMASH with MALDI-TOF MS analysis, we revealed that the antifungal activity was due to the production of the lipopeptides herbicolins A and B.

The antifungal compounds produced by the strain indicated potent activity against sixteen of the seventeen tested fungal indicators. Specifically, the purified lipopeptides were active against the food spoilage molds *A. niger* UCC, *A. persicinum* DSM 1052, *A. alternata* DSM 1102, *F. solani* DSM 1164, *S. brevicaulis* DSM 1218, *G. candidum* KH-1730, *Geotrichum* sp. UCC, *P. variotii* UCC, *Cladosporium* sp. UCC, *B. nivea* UCC, P. *notatum* UCC, and Phoma sp. UCC, and the yeasts *C. parapsilosis* UCC, *Y. lypolitica* strain 78-003, *S. cerevisiae* type strain Sa-07140, and *Z. bailii* strain Sa-1403. The strongest activity was observed against the yeast *S. cerevisiae* type strain Sa-07140, *Z. bailii* strain Sa-1403, and *Y. lipolytica* strain 78-003 (MIC 0.2 μg/ml). Our results are in agreement with Xu and coworkers (Xu et al., 2022), who revealed strong inhibitory activity against *S. cerevisiae* BY4741 (EC50 = 0.3 μg/ml). The broad spectrum antifungal activity of herbicolin A is supported by previous studies, indicating that the lipopeptide can prevent the growth of several plant-pathogenic fungi, such as *Schizonella melanogramma, Botrytis cinerea, Urocystis occulta* (Winkelmann et al., 1980), *Fusarium culmorum, Puccinia recondita* (Kempf, 1989), *Fusarium graminearum* (Xu et al., 2022), *Verticillium dahliae, Helminthosporium sativum, Pyrenophora graminea, Cladosporium* sp., *Mycosphaerella graminícola, Phialophora fastigiata, Gaeumannomyces graminis* var. *tritici, Colletotrichum coccodes, Fusarium culmorum, Verticillium chlamydosporum, Botrytis allii, Botrytis cinerea, Botrytis fabae, Monilinia fructigena, Rhizoctonia solani, Rhizoctonia cerealis and Rhizoctonia tuliparum* (Matilla et al., 2023). According to Xu and coworkers (Xu et al., 2022), the antifungal activity of herbicolin A is based on interactions with ergosterol detected in the cell membranes, leading to the disruption of ergosterol-containing lipid rafts. The fact that ergosterol is a basic component of fungal cell membranes (Rodrigues, 2018) emphasizes the efficiency of these components for anti-fungal applications.

Whole genome sequencing analysis and genome assembly revealed that the gene cluster encoding for herbicolins A and B is plasmid-encoded. The presence of a tyrosine-type recombinase/integrase and an Rpn recombination-promoting transposase adjacent to the gene cluster suggests that the gene cluster could be subject to horizontal transfer. Genome mining revealed a complex non-ribosomal biosynthesis and indicated low sequence similarity (6%) with structural components of the gene clusters encoding for hassallidin C (Vestola et al., 2014). Hassallidins are known antifungal glycolipopeptides which are widespread among cyanobacteria. Comparison with NRPS produced by other *P. agglomerans* strains showed high sequence similarity (≥ 98%) with the structural components of the plasmid-encoded gene cluster for herbicolins A and B, previously described by Matilla and coworkers, and Xu and coworkers (Matilla et al., 2023; Xu et al., 2022). Interestingly, from the 284 *P. agglomerans* genomes that have been so far submitted to the NCBI database (https://www.ncbi.nlm.nih.gov/), only *P. agglomerans* ZJU23 and *P. agglomerans* 9Rz4 were identified to contain this gene cluster. Also, the two strains originate from geographically distant locations. Specifically, *P. agglomerans* ZJU23 was isolated from *Fusarium* fruiting body microbiome in China, whereas *P. agglomerans* 9Rz4 was from the oilseed rape rhizosphere in Germany. This indicates that the gene cluster encoding for herbicolins A and B is not widespread among *P. agglomerans* strains. In addition, Matilla and coworkers (Matilla et al., 2023), suggested that the herbicolin A gene cluster was obtained by horizontal gene transfer, due to the presence of remnants of transposases, and its higher G + C content compared to the rest of the plasmid.

Some differences that were identified in our gene cluster were the lack of two open reading frames (*acbA* and *acbB*) and three open reading frames (*hbcA* to *hbcC*) compared to the biosynthetic clusters encoding for ZJU13 and 9Rz4, respectively. Domain identification analysis indicated the lack of one adenylation subunit compared to HbcD from *P. agglomerans* ZJU23 and *P. agglomerans* 9Rz4, and one condensation domain compared to HbcD from *P. agglomerans* 9Rz4. Searching for additional genes encoding adenylation domain containing proteins located around the putative herbicolin A gene cluster in *P. agglomerans* APC 4211 identified only a single adenylation domain predicted to encode anthranilic acid. Increased nucleotide similarity (≥ 98%) was also observed for genes encoding for a 3-oxoacyl-(acyl-carrier-protein) reductase, an MbtH protein, and a glycosyltransferase, suggesting conserved post-translation modifications reactions during herbicolins A and B production. The gene cluster also encoded a putative transporter and a multidrug export protein possibly involved in the export of the lipopeptides to the extracellular environment. Furthermore, an *abi* family immunity gene, which possibly encodes self-immunity to the strain was also detected.

To enhance the production of herbicolins A and B, fermentation conditions for *P. agglomerans* APC 4211 were optimized by assessing different growth conditions (medium composition and incubation time). Maximum lipopeptides production was achieved in LM17T medium following 48 hours of incubation. The optimization experiments of Wang and coworkers ^52^ suggested that supplementation of King’s B medium (King et al., 1954) with corn steep liquor, glycerol, CaCl_2,_ and threonine contributed to the highest herbicolin A production. In our experiments, glycerol did not contribute to increasing the fermentation yield. In agreement with our results, increasing the incubation time had a positive effect on herbicolin A production. To further improve herbicolins A and B isolation, we applied the freeze and squeeze technique. This approach contributed to a drastic increase in the size of the inhibition zone against all the tested indicators, including *S. cerevisiae* type strain Sa-07140, *Y. lipolytica* strain 78-003, and *Geotrichum* sp. UCC, for which no inhibition activity was previously observed using the well diffusion assay.

To explore whether the detected NRPs had theoretical masses corresponding to the lipopeptides herbicolin A and herbicolin B, HPLC fractionation was performed followed by MALDI-TOF MS analysis. This analysis revealed ions with masses consistent with herbicolin A (1300.8 Da), a peptide with an additional 18 Da mass (1300.8 Da +18 Da), and the putative mass of herbicolin B (1138 Da). The masses detected in the colony mass spectrum correlated well with the masses detected from the HPLC-purified antimicrobials, indicating that herbicolins A and B are the major compounds produced by the strain. Evaluation of the MICs of purified herbicolin A, purified herbicolin B and combination of herbicolin A and herbicolin B, as produced by *P. agglomerans* APC 4211, indicated low MICs against all tested indicators, with the lowest MIC to be indicator-specific. Furthermore, the activity of the partially-purified CFS was unaffected by proteinase K and trypsin and remained stable after heat treatment (from 40 °C to 100 °C for 15 minutes), highlighting the stability of the lipopeptide. These characteristics are considered important for possible biotechnological applications of the herbicolins A and B.

To evaluate the efficacy of herbicolins A as natural biopreservative, compared to natamycin, a natural preservative with generally recognized as safe (GRAS) status produced by *Streptomyces natalensis* (Meena et al., 2021), as well as other anti-fungal antibiotics that exist in the market, including fluconazole, ketoconazole, itraconazole, and terbinafine, we tested its MIC against fungal indicators. Herbicolin A was shown to be more effective compared to the other antibiotics against *Geotrichum* sp. UCC, and *S. brevicaulis* DSM 1218. This is specifically important because *Geotrichum candidum* is among the major creamy dairy products spoilage fungi, leading to severe economic consequences for the dairy products industry (Kamilari et al., 2023). Also, *S. brevicaulis* was associated with human infections, including eye infections (keratitis and endophthalmitis), onychomycosis, disseminated skin lesions, pneumonia, and other infections in immunocompromised patients (Richardson and Lass-Flörl, 2008). It is noteworthy that some reports indicate that treatments with terbinafine, itraconazole, ketoconazole, and fluconazole may lead to side effects, including allergic reactions, stomach pain, diarrhoea, or headache, and in more severe cases hepatotoxicity, or breathing difficulties (Gupta et al., 2023). Testing of chemical preservatives revealed that only sorbic acid was effective against the tested indicators but with higher MIC values compared to herbicolin A (0.187 mg/ml to 0.75 mg/ml). Excessive intake of sorbic acid may have harmful consequences for human health, such as carcinogenic and mutagenic effects (Samoylov et al., 2021). Furthermore, herbicolins A and B were not cytotoxic against human epithelial cells at a concentration of 50 μg/ml. These results provide an indication of the safety of herbicolins A and B, but more extensive studies need to be performed before to determine their safey for biopreservative applications.

Considering that milk and milk derivatives are essential components in human diets, and among the highest affected and wasted foodstuffs annually due to fungal contamination (Martin et al., 2021), we evaluated the anti-fungal effects of herbicolin A against *A. niger* UCC, *P. notatum* UCC, and *Geotrichum* sp. UCC using skim milk as a dairy food model. We showed that the concentration of 5 μg/ml eliminated the presence of *A. niger* UCC and *P. notatum* UCC and reduced significantly the concentration of *Geotrichum* sp. UCC after 3 days of incubation at 30 °C. A significant reduction in the concentration of these fungi was observed following 2.5 μg/ml of lipopeptides administration. These results demonstrate that herbicolin A was able to maintain activity against the tested fungal indicators when applied as a biocontrol agent in milk. However, the economic ramifications of adding the antimicrobial in these amounts has yet to be determined. Overall, this study highlights the potency and efficiency of herbicolin A as potential candidates for biological control of food ecosystems and consequently may assist in improving food safety.

## 5. Conclusion

Food spoilage due to fungal contamination represents a critical challenge for the food industry. The increasing consumer demand for natural, free-from-synthetic additives preservatives food products stresses the necessity for the identification of natural, effective, and safe biopreservatives against spoilage and pathogenic fungi. To our knowledge, this is the first study providing evidence of the efficient and safe administration of herbicolin A, produced by the strain *P. agglomerans* APC 4211, as biocontrol agents for food ecosystems. Still, more tests and clinical trials need to be performed to prove their biosafety and economic viability. Additionally, the rapid increase of fungi developing genetic changes to survive drugs highlights the importance of discovering and testing NRPs as a possible solution to this global threat.

## Author statements

### Author contributions

Conceptualization, J.R., C.S., R.P.R. and C.H; methodology, E.K., P.M. O’C, M.A.S.N; data collection: E.K., software, E.K., validation, E.K.; formal analysis, E.K.; investigation, E.K.; resources, C.S. and R.P.R., C.H; data curation, E.K.; writing—original draft preparation, E.K.; writing—review and editing, J.R., C.S., R.P.R., C.H, P.D; visualization, E.K.; supervision, C.S., R.P.R., J.R., C.H; project administration, C.S., R.P.R., C.H; funding acquisition, R.P.R., C.S., C.H, O.F, J.W, D.H, A.D. All authors have read and agreed to the published version of the manuscript.

## Conflicts of interest

All authors declare no financial or non-financial competing interests

## Funding information

This research was funded by the Science Foundation Ireland (SFI) through APC Microbiome Ireland, under Grant number SFI/12/RC/2273, by Kraft-Heinz company funding, and by the European Union (ERC, BACtheWINNER, Project No. 101054719). However, the views and opinions expressed are those of the author(s) only and do not necessarily reflect those of the European Union or the European Research Council Executive Agency.

## Ethical approval

Not applicable

## Consent for publication

Not applicable

## Acknowledgements

We would like to thank Ruth Fair from Bandon Grammar School, Co. Cork, Ireland for her contribution in sample collection and colonies isolation.

## Data availability

The datasets generated and/or analysed during the current study are available in NCBI repository. https://submit.ncbi.nlm.nih.gov/subs/wgs/SUB15169161/overview

## Notes

### Competing Interest Statement

The authors have declared no competing interest.

