## Supplementary material for "Herbicolin A, an antifungal lipopeptide produced by *Pantoea agglomerans* APC 4211 is a promising biocontrol agent against food spoilage fungi"

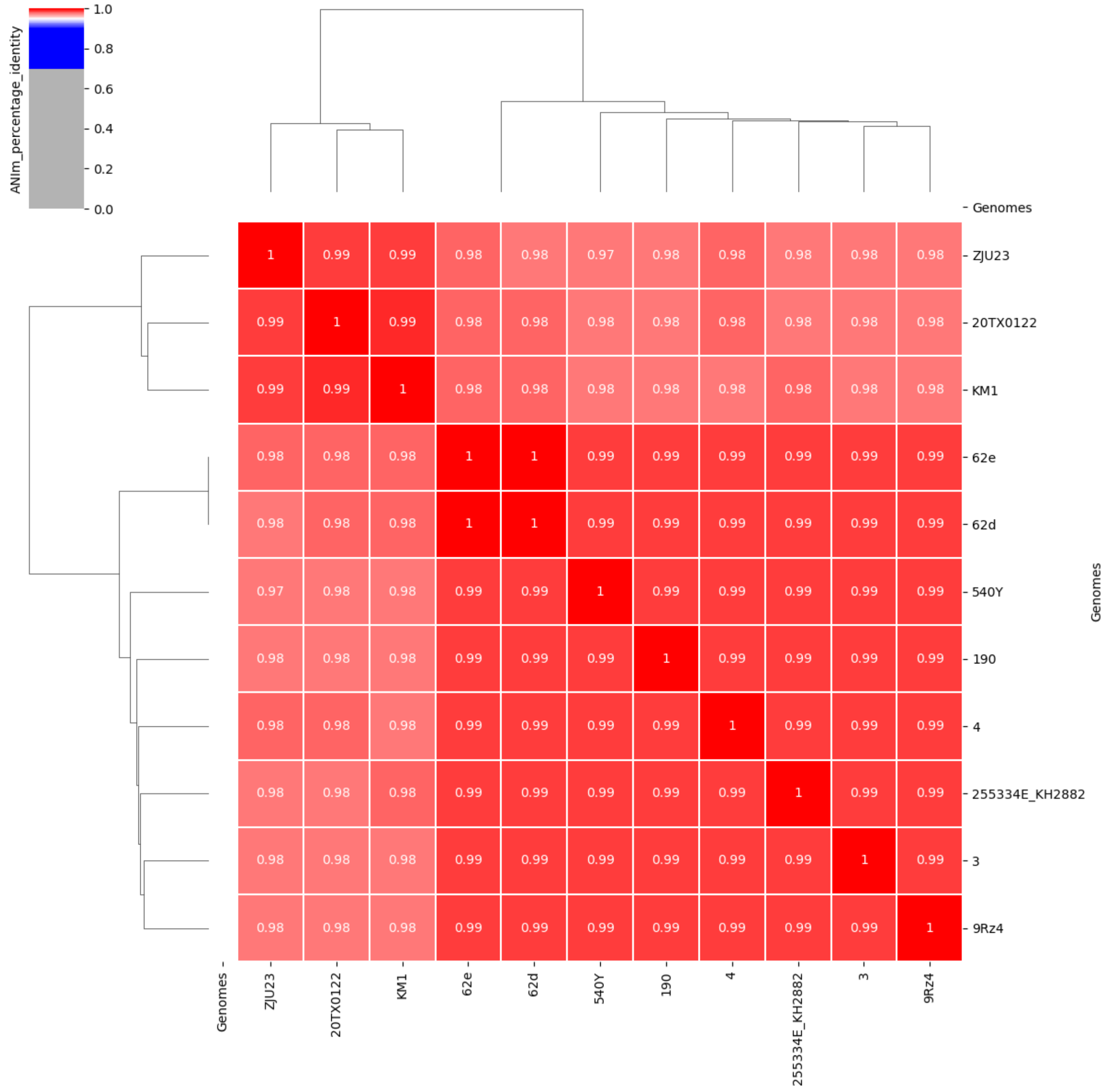


**Supplementary Figure 1**. Comparison of the whole genome of *P. agglomerans* APC 4211 against other 10 *P. agglomerans* complete circular genomes

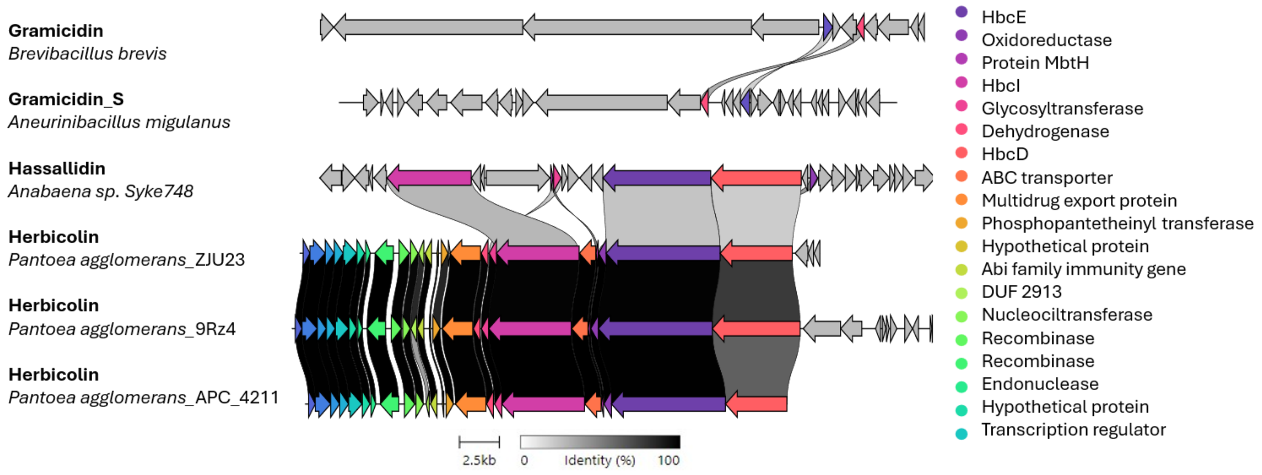


**Supplementary Figure 2**. Schematic representation of the similarity among the gene clusters associated with herbicolin A from *P. agglomerans* ZJU23 and *P. agglomerans* 9Rz4, hassallidin from *Anabaena sp. Syke748*, gramicidin from *Brevibacillus brevis*, and gramicidin S from *Aneurinibacillus migulanus,* and the identified gene cluster detected in *P. agglomerans* APC 4211 genome. Genes are color-coded according to the putative role of each protein.

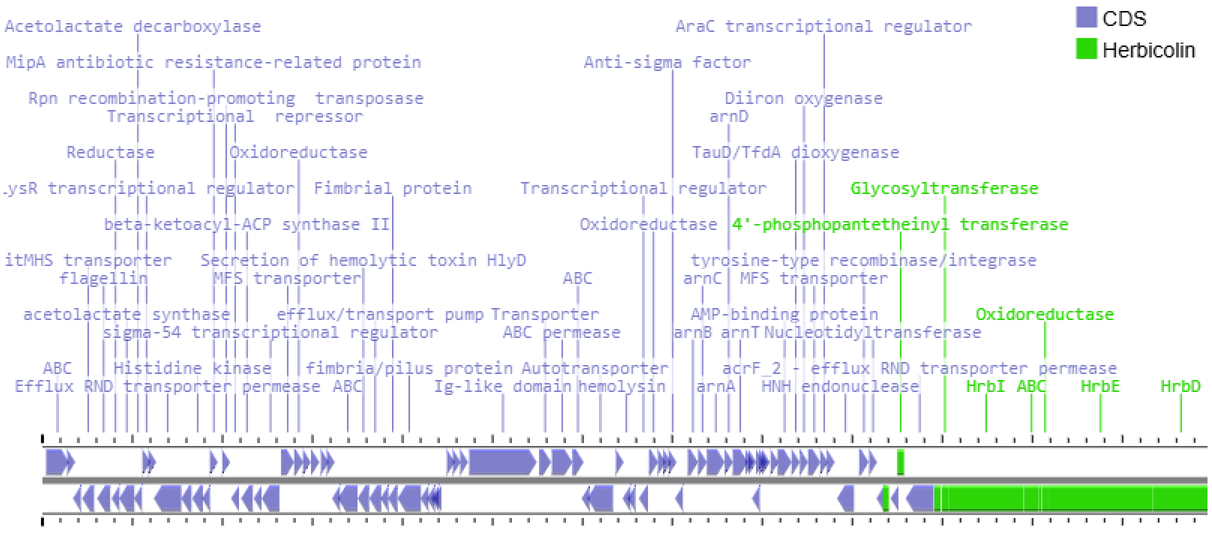


**Supplementary Figure 3.** Schematic representation of the genes detected in the plasmid encoding for herbicolins A and B.

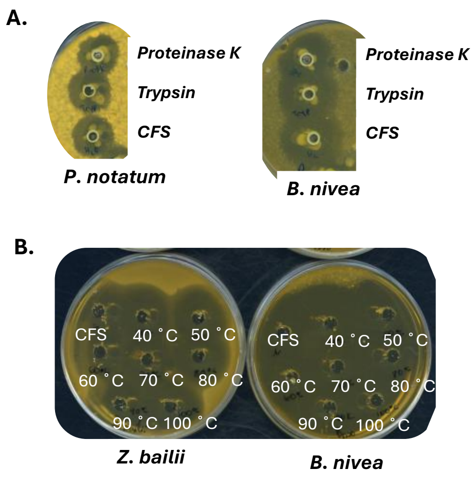


**Supplementary Figure 4**. Sensitivity of the partially purified CFS to proteinase K and trypsin using the well diffusion assay, with *P. notatum* UCC and *B. nivea* UCC as indicator strains. **B.** Stability of the partially purified CFS to exposure to different temperatures (40 ˚C to 100 ˚C) for 15 minutes, using the well diffusion assay, against *Z. bailii* strain Sa-1403 and *B. nivea* UCC.

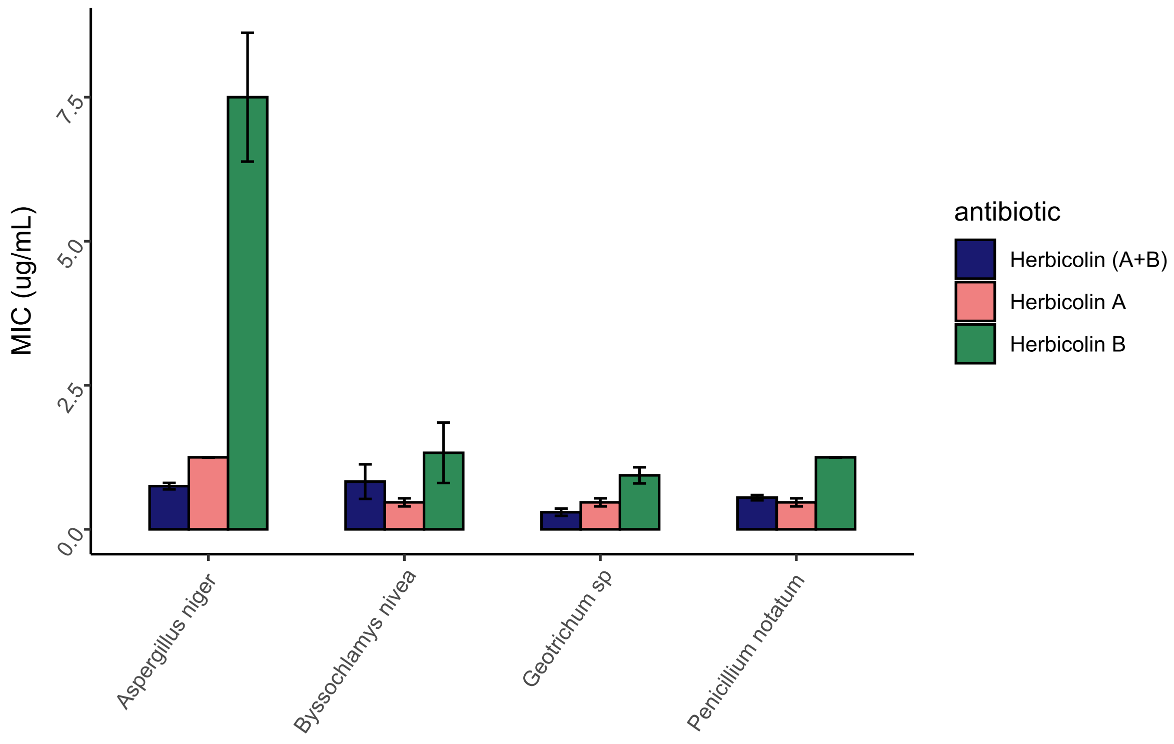


**Supplementary Figure 5.** Comparison of the activity of herbicolins A and B, compared to purified herbicolin A and purified herbicolin B.

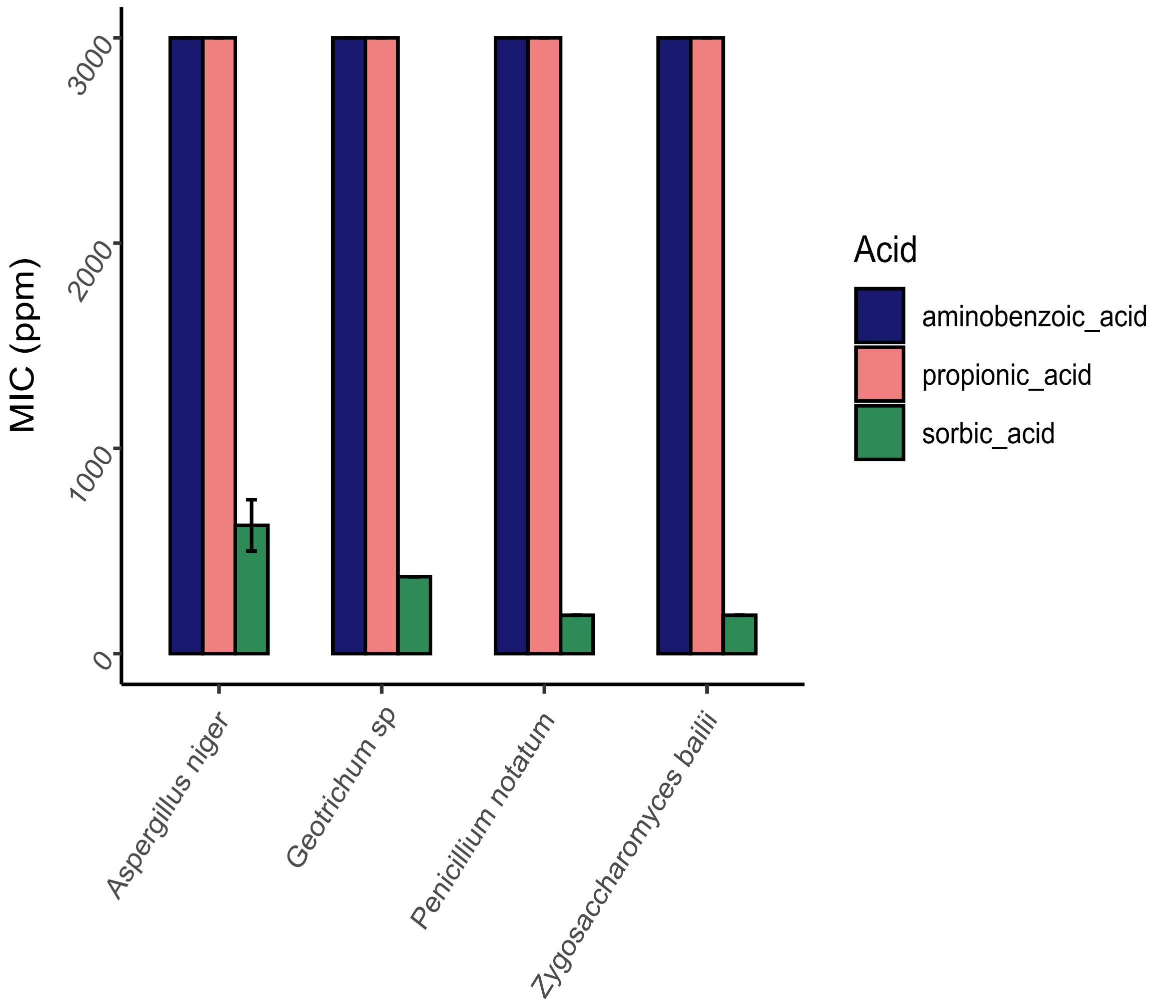


**Supplementary Figure 6.** Evaluation of fungal indicators MIC values against 4-aminobenzoic acid, propionic acid, and sorbic acid.
